# A Whole-Cell Screening Platform to Discover Cell Adhesion Molecules that Enable Programmable Bacterial Cell-Cell Adhesion

**DOI:** 10.1101/2023.12.03.569830

**Authors:** Po-Yin Chen, Yung-Chih Chen, Po-Pang Chen, Kuan-Ting Lin, Wei-Le Wang, Kuo-Chiang Hsia, See-Yeun Ting

## Abstract

Developing programmable bacterial cell-cell adhesion is of significant interest due to its versatile applications. Current methods that rely on presenting cell adhesion molecules (CAMs) on bacterial surfaces are limited by the lack of a generalizable strategy to identify such molecules targeting bacterial membrane proteins in their natural states. Here, we introduce a whole-cell screening platform designed to discover CAMs targeting bacterial membrane proteins within a synthetic bacteria-displayed nanobody library. Leveraging the potency of the bacterial type IV secretion system—a contact-dependent DNA delivery nanomachine—we have established a positive feedback mechanism to selectively enrich for bacteria displaying nanobodies that target antigen-expressing cells. Our platform successfully identified functional CAMs capable of recognizing three distinct outer membrane proteins (TraN, OmpA, OmpC), demonstrating its efficacy in CAM discovery. This approach holds promise for engineering bacterial cell-cell adhesion, such as targeted antimicrobial interventions in the microbiome by deploying programmed inhibitor cells.

## INTRODUCTION

Cell-cell interactions are crucial in shaping the intricate structures of microbiomes and facilitating essential functions, including genetic exchange, metabolite transfer, and cellular stress responses. The ability to engineer and manipulate such multicellular arrangements not only enhances our understanding of microbial communities, but also serves as a basis for developing synthetic platforms, which can enable the development of functional microbial consortia with diverse biotechnological applications^3, 4^. For example, multicellular bioconversion represents an area of significant scientific interest, whereby spatially organized microbial communities efficiently convert complex substrates into valuable products^5, 6^. Moreover, drawing inspiration from natural biofilms, self-repairing living biomaterials with cell-cell adhesion properties show regenerative capabilities and display improved resilience^5, 7^. Furthermore, engineered intercellular adhesion could have profound impacts on targeted manipulations within complex microbial populations, enabling precise and efficient interventions against specific bacteria^8-10^.

Recent progress in the field of directed cell-cell adhesion has involved the use of bacteria displaying synthetic cell adhesion molecules (CAMs), such as nanobody-antigen or coiled-coil peptide pairs, to facilitate intercellular binding^11, 12^. These strategies are a rational approach for programming multi-microbial materials. However, despite their potential, certain limitations persist due to restrictions in the number of effective binding pairs that can be deployed for cell-cell adhesion purposes. The current development bottleneck arises from using conventional methods to identify potential binders, such as phage- or yeast-display selection, which were not originally designed to identify functional CAMs for bacteria. These methods often present challenges, including 1) poor expression or insufficient display of selected molecules on bacterial surfaces; and 2) reduced binding affinity due to interference from bacterial surface components, such as other membrane proteins or extracellular polysaccharides^8, 9^. Consequently, the laborious process of testing functional CAMs for suitable display on bacterial surfaces is a primary obstacle in the design of engineered microbial communities^11, 12^.

To address this challenge, we sought to develop a high-throughput screening platform for the identification of functional CAMs with the following properties: *i*) stable display on the bacterial surface; *ii*) targeting of surface antigens in their native conformations; and *iii*) the ability to mediate intercellular adhesions between bacteria. A whole-cell-based platform has the potential to meet these requirements by screening surface-displayed CAMs against bacterial cells harboring target membrane protein antigens. Similar strategies have been implemented to isolate binders against mammalian membrane proteins in their native forms^13,14^. However, analogous platforms for targeting bacterial membrane proteins are limited.

In this study, we have developed a directed evolution workflow based on a whole-cell screening approach, aimed at enriching for CAM-displaying cells by using antigen-expressing bacteria as bait. To avoid spontaneous cell aggregates derived from non-specific intercellular binding, our workflow incorporates a positive feedback mechanism at each panning stage, exploiting the contact-dependent DNA delivery nanomachine that operates between interacting bacteria, *i.e.,* the type IV secretion system (T4SS) (**Fig. 1a**). T4SS is a widespread pathway that naturally conjugates genetic material from donor to recipient cells through direct cell-cell interactions. This secretion system is known to target cells with low specificity, unless a strong and stable mating junction forms between bacteria^15^. Elegant works by Robledo *et al*. and Li *et al.* demonstrated that such mating pairs could be achieved by synthetic cell-cell adhesions, promoting target bacterial conjugation even under fluid conditions when cell-cell contacts are transient^9, 10^. Therefore, we posited that the selective properties of T4SS, mediated by synthetic CAMs, could effectively facilitate conjugative transfer of desired genes, such as antibiotic-resistance markers, allowing transconjugants to survive on selective media during the selection process. We anticipated that this approach would result in a positive enrichment for bacteria displaying cognate CAMs from synthetic nanobody libraries (**Fig. 1b**).

**Fig. 1.**
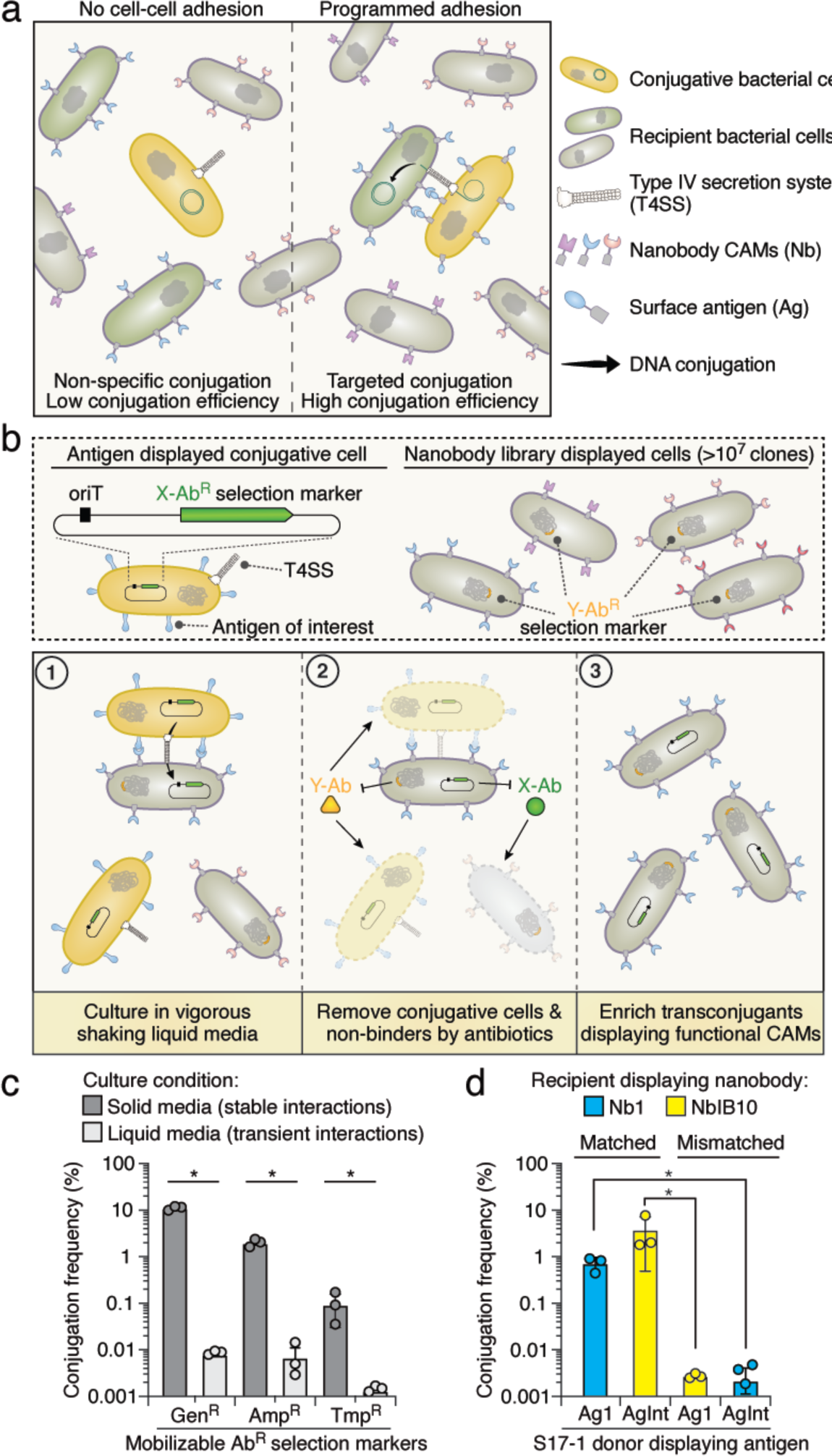
Establishment of the whole-cell-based CAM discovery platform in liquid medium. (**a**) Schematic of the strategy for facilitating precise DNA conjugation between specific cells through synthetic cell-cell adhesion. Bacteria expressing diverse nanobodies (Nb) and conjugative cells with surface antigens (Ag) are involved. (**b**) The whole-cell-based selection platform is designed to identify nanobody CAMs targeting specific antigens of interest. Conjugative cells expressing the target antigen and cells displaying the nanobody library are employed. ① Within a liquid culture, contact-dependent DNA conjugation is enhanced through strong binding between cognate nanobodies and antigen pairs (both labeled in blue). ② After conjugation, antibiotics are used to eliminate conjugative cells and recipients expressing non-nanobody binders. ③ After a single round of selection, transconjugants carrying nanobodies with a relatively strong binding affinity toward the target antigens are enriched. (**c**) Conjugation frequencies (transconjugants/total recipients) of conjugative plasmids (pGenR, pAmpR, and pTmpR) were assessed under both solid and liquid growth conditions for a 6-hour duration. *E. coli* S17-1 serves as the conjugative donor strain, and MG1655 serves as the recipient strain. (**d**) Conjugation frequency of pGenR under the liquid growth condition. Recipient cells expressing nanobodies were co-cultured with donor cells displaying cognate or orthogonal antigens. Data are presented as means ± SD. n = 3 technical replicates, representative of two biological replicates. *p < 0.05, t-test. See also **Supplementary Fig. 1**.

Here, we demonstrate the application of our whole-cell screening platform to identify functional CAMs that recognize three distinct surface-localized proteins—TraN, OmpA, and OmpC—in their native conformations. We show that mounting these nanobody CAMs on bacterial surfaces not only induces pronounced bacterial cell-cell adhesion, but also enables targeted cell depletion in the microbial community by using programmed inhibitor cells equipped with the nanobodies^8^. By providing a means to engineer tailored interactions between bacteria, our platform opens up new possibilities for the development of advanced synthetic applications in basic research, biotechnology, and clinical settings.

## RESULTS

### Facilitating Selective Gene Transfer in Mixed Populations through Synthetic Cell-Cell Adhesion

We began by establishing a selection platform aimed at achieving targeted gene transfers within mixed bacterial populations. This platform relied on the utilization of conjugative donor cells equipped with surface-exposed antigens that could distinguish recipient cells displaying cognate binders from those displaying non-cognate controls. This discerning recognition enabled selective gene transfer events within the mixed population, as depicted in **Figure 1a**. For proof-of-concept experiments, we employed characterized antigen (Ag) - nanobody (Nb) pairs in conjunction with a previously developed intimin display system^11, 16^.

Our initial assessments involved testing the feasibility of this strategy using *Escherichia coli* S17-1 strain (ATCC 47055), a genetically manipulable bacterium with active conjugation ability under standard laboratory conditions. Consistent with previous findings, S17-1 with rigid pili inefficiently transferred antibiotic-resistance markers on mobilizable genetic elements to susceptible recipients in liquid media (**Fig. 1c**), where cell-cell interactions are transient^17,18^. However, the conjugation frequency was significantly enhanced (100- to 1000-fold) when intercellular contacts were enforced by bacterial growth on solid agar plates. The enhancement in conjugation frequency was not observed for an S17-1 strain with a dysfunctional T4SS (Δ*trbE*) for both liquid and solid conditions^19^ (**Supplementary Fig. 1a**).

Next, we introduced antigen and nanobody expression systems into the conjugative donor and recipient strains, respectively. To enable comparative analyses, we incorporated two distinct characterized cell adhesion binding pairs, namely Ag1-Nb1 and AgInt-NbIB10^11, 20^. Conjugation frequencies of S17-1 were markedly increased in liquid cultures when recipient cells displayed cognate nanobodies, whereas only mild effects were detected for non-cognate nanobodies (**Fig. 1d**). Importantly, this selective gene transfer phenomenon was contingent on T4SS being functional within the S17-1 strain (**Supplementary Fig. 1b**). These results support the notion that Ag-Nb-mediated cell-cell adhesion plays an important role in directing the T4SS toward recipient cells when cell-cell interactions are unstable in liquid culture.

### Iterative Selection Enables Discriminative Enrichment for Low-Abundance Bacteria Displaying Cognate Nanobodies

A hallmark of our devised workflow is its capacity to differentially interact with target and non-target cells. To explore this feature further, we subjected antigen (Ag1)-displaying conjugative bacteria to recipient cells expressing matched nanobodies (Nb1) across varying degrees of dilution in the presence of cells expressing a non-matched control. Under these conditions, we detected an enhanced conjugation frequency to recipients when control cells outnumbered target cells by factors of 10-, 100-, and 1,000-fold (**Fig. 2a**). The enrichment for cells expressing matched nanobodies (Nb1) was further verified by diagnostic PCR and sequencing (**Supplementary Fig. 2a**,b). However, we noted that when the target cells were diluted 10,000-fold in the initial pool, no statistically significant difference in conjugation frequency was observed, indicating an upper limit for this discriminant capability (**Fig. 2a**).

**Fig. 2.**
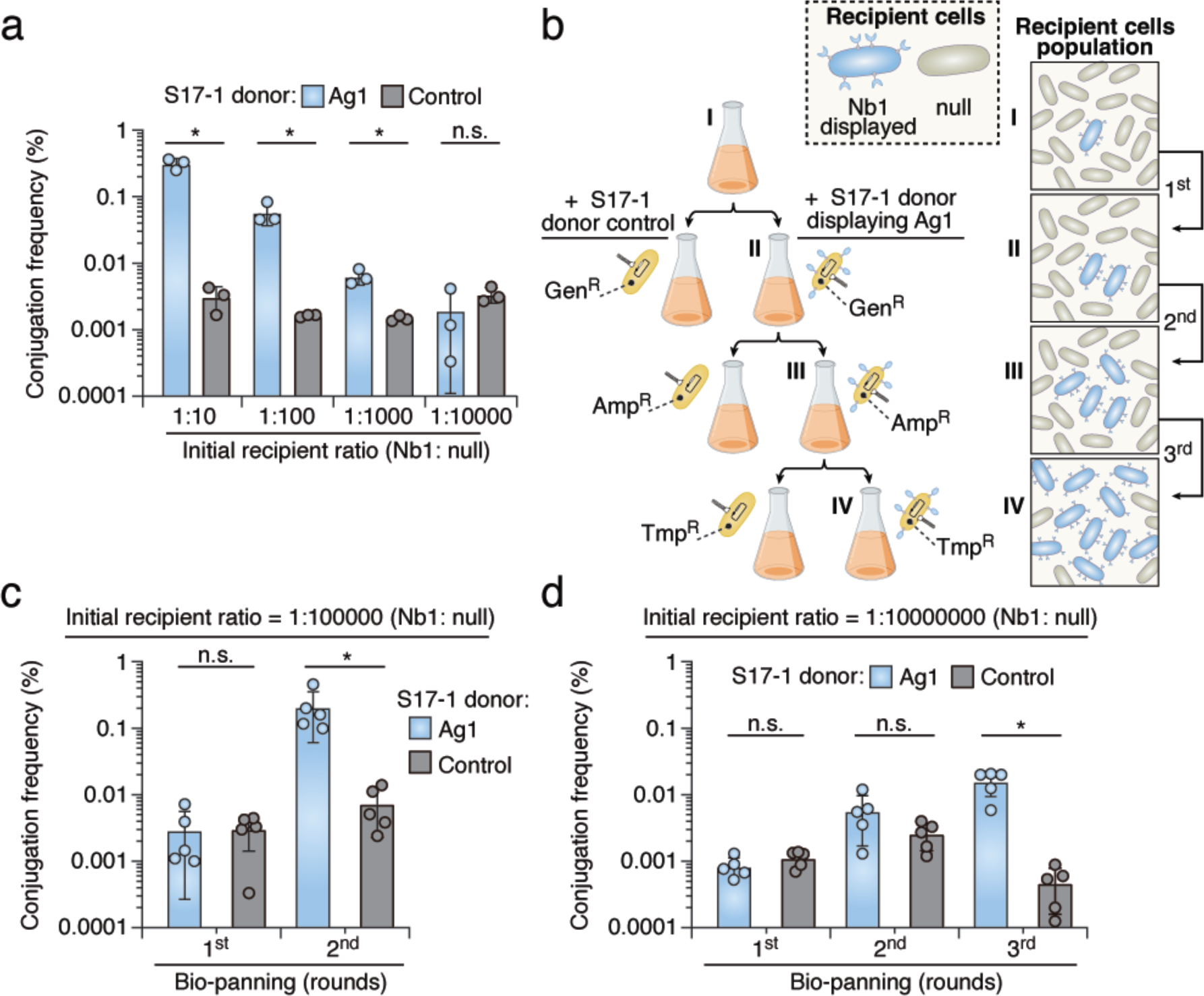
Iterative rounds of bio-panning enhance the discriminant capacity of the CAM selection platform. (**a**) Conjugation frequency of pGenR in conjugation assays within liquid culture between recipient and donor displaying Ag1 or Control (without antigen displayed). The recipient culture expressing Nb1 was 10-, 100-, 1000-, and 10000-fold diluted with the culture expressing non-matched null cells. (**b**) Schematic of three-round selection. For each round, donors (yellow) carry plasmids containing antibiotic-resistance genes against gentamicin (GenR), ampicillin (AmpR), or trimethoprim (TmpR) as distinct selection markers. The cartoon illustration on the right represents the progressive increase in the abundance of recipient cells displaying Nb1 (blue) in a mixed population with non-matched null cells (grey) during the selection process. (**c**) Conjugation frequency in two rounds of bio-panning. The initial population of recipient cells expressing Nb1 and null cells were mixed at a ratio of 1:10^5^. (**d**) Conjugation frequency in three rounds of bio-panning. The initial population of recipient cells expressing Nb1 and null cells were mixed at a ratio of 1:10^7^. Data are presented as means ± SD. n ≥ 3 technical replicates, representative of two biological replicates. *p < 0.05, t-test. See also **Supplementary Fig. 2**.

Central to our selection strategy is the screening of a library comprising a minimum of 10^7^ distinct nanobody clones. Given this objective, we postulated that iterative rounds of bio-panning could overcome the dilution threshold. Therefore, we harvested transconjugants from the first round of selection and introduced them to a new round of conjugation and selection, using distinct mobilizable antibiotic-resistance markers (**Fig. 2b**). After two rounds of selection, transconjugants having matched nanobody (Nb1) were enriched from a pool of cells initially mixed at a ratio of 1:10^5^ (Nb1:null) (**Fig. 2c and Supplementary Fig. 2c**), and three rounds of bio-panning resulted in successful isolation from a population of cells initially mixed at a ratio of 1:10^7^ (Nb1:null) (**Fig. 2d and Supplementary Fig. 2d**). These outcomes demonstrate the capacity of our designed workflow to exhibit notable specificity towards low-abundance target cells within mixed bacterial populations.

### Discovery of a Nanobody Targeting the Natural Adhesin TraN

Building on the success of our selection workflow, next we sought to determine if our approach could yield nanobody clones capable of targeting protein antigens naturally present on bacterial cell surfaces. We were particularly interested in TraN, a bacterial outer-membrane protein with accessible epitopes on the cell surface that is involved in stabilizing mating pairs and facilitating horizontal gene transfers^15, 21^. Most relevant to our study, TraN functions as a natural adhesin, promoting cell-cell adhesion upon binding to its specific receptor. This property renders it an ideal module for enabling programmable cell-cell adhesion and represents an appropriate choice for our proof-of-principle experiments.

Accordingly, we assessed the feasibility of isolating bacteria expressing cognate nanobodies from a mixed population by displaying the TraN antigen on S17-1 conjugative cells. Our experimental workflow encompassed incubating of S17-1 cells with a bacteria-displayed nanobody library derived from a synthetic pool comprising approximately 10^7^ clones of distinct nanobodies^22^. Following each round of selection, we harvested colonies and conducted high-throughput sequencing to confirm enrichment. Our analyses revealed a progressive increase in the abundance of specific nanobody clones within the library pool across successive rounds of selection (**Fig. 3a and Supplementary Table 3**). To validate TraN-dependent cell-cell adhesion, we selected the top three unique sequences from the final round of panning, introduced them to model *E. coli*, and conducted a macroscopic cell aggregation assay using optical density (OD_600_) measurements to quantify the binding strength and specificity between cells^11^. If the nanobodies possessed CAM capacity, we expected to observe rapid precipitation in a cell mixture of the corresponding nanobody- and antigen-displaying strains. Consistent with our screening results, we observed a significant reduction in supernatant OD_600_ in a mixture of cells expressing one of the top nanobodies and cells displaying TraN, indicating that the specific nanobody on the cell surface served as a functional CAM in binding to TraN and precipitating cells from culture (**Fig. 3b and Supplementary Fig. 3a**). This result prompted us to pursue an in-depth characterization of the isolated TraN-specific nanobody, hereafter referred to as Nb^traN^ (**Fig. 3c**).

**Fig. 3.**
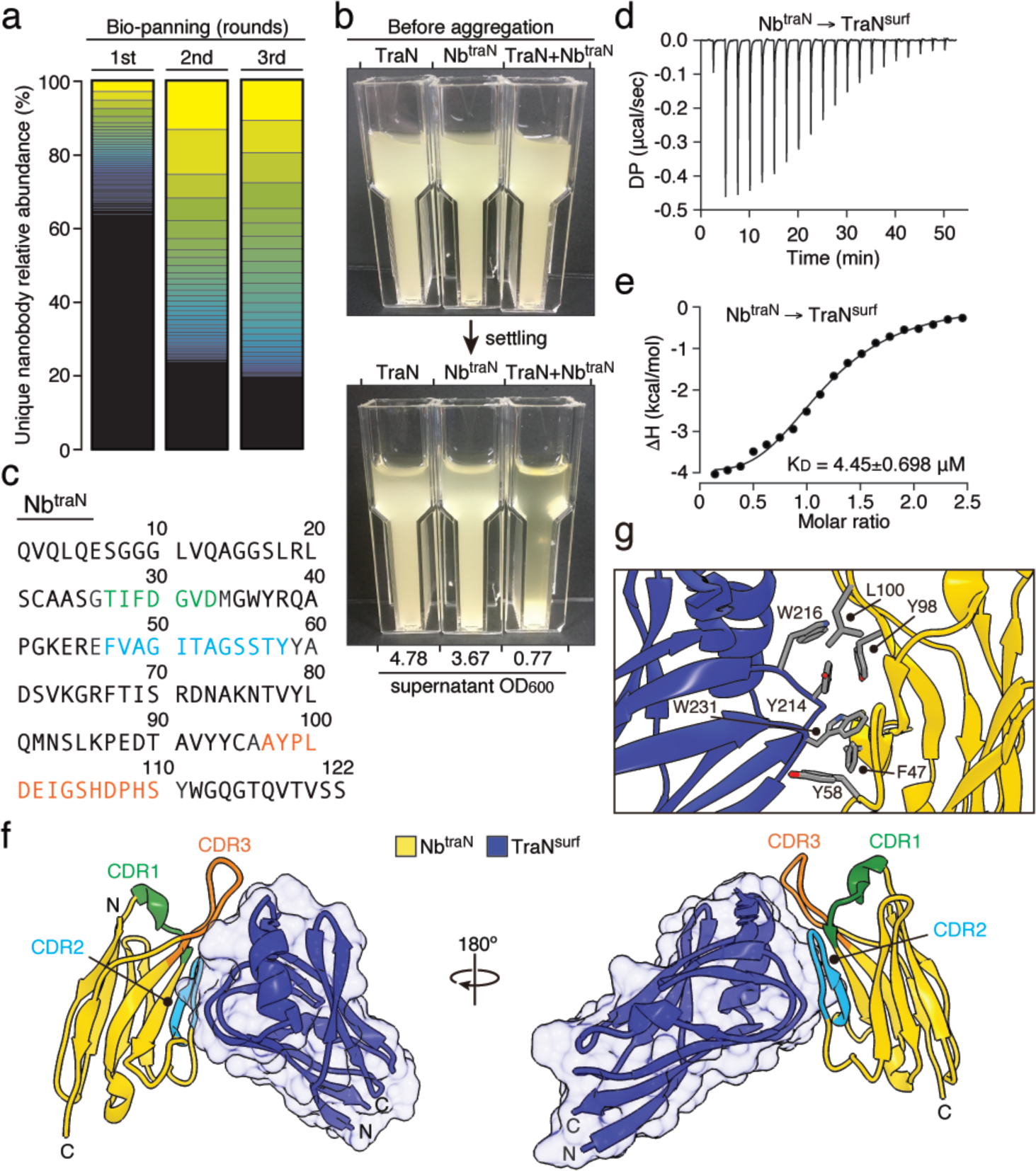
Identification of nanobody recognizing TraN via the CAM discovery platform. (**a**) Stacked bar plot illustrating the changes in the relative abundance of individual nanobodies within the population across three rounds of bio-panning to enrich for bacteria displaying TraN-binding nanobodies. Sequences were discerned through the CDR regions of the nanobody gene. The relative abundances of all unique nanobodies identified are shown, and the respective counts are detailed in **Supplementary Table 3**. Colors denote the ranking of nanobody abundance from the highest to lowest. (**b**) Macroscopic aggregation analysis to assess cell-cell adhesion. Cultures of *E. coli* strains displaying TraN or Nb^traN^ were incubated either individually or in combination. The OD_600_ of the supernatant after the settling period is shown below the image. (**c**) The amino acid sequence of Nb^traN^, shows CDR1, CDR2, and CDR3 in green, blue, and orange, respectively. (**d, e**) The ITC binding profiles of Nb^traN^ with TraN^surf^ at 25 °C. (**f**) Crystal structure of Nb^traN^ (yellow) in complex with TraN^surf^ (blue) (PDB: 8X7N). CDR1, CDR2, and CDR3 are presented in green, blue, and orange, respectively. (**g**) A magnified view of the interacting residues is indicated between Nb^traN^ and TraN^surf^. See also **Supplementary Table 1**.

As expected, we detected a stable interaction between purified Nb^traN^ and the surface-exposed domain of TraN (TraN^surf^) via size-exclusion chromatography (**Supplementary Fig. 3b**,c). Further analysis revealed a binding affinity (K_D_) of 4.45 μM between Nb^traN^ and TraN^surf^ (**Fig. 3d,e**), confirming the specificity of the interaction. To gain insights into the interaction between Nb^traN^ and TraN, as well as how it promotes cell-cell adhesion in bacterial mixtures, we determined a 3.67 Å X-ray crystal structure of Nb^traN^ in complex with TraN^surf^ (PDB: 8X7N) (**Fig. 3f,g and Supplementary Table 1**). The Nb^traN^/TraN^surf^ complex structure was determined by molecular replacement using nanobody homolog, Nb.b201, as a search model, which has an identical fixed framework region to that of Nb^traN^ (PDB: 5VNV)^22^. In our structure, Nb^traN^ and TraN^surf^ share an extensive interaction surface consisting of two contact regions that are primarily formed by the complementarity-determining regions 2 and 3 of Nb^traN^ (CDR2, CDR3) (**Fig. 3f**). In contrast, CDR1, the other variable loop in Nb^traN^, did not present any direct interaction with TraN^surf^ in our structure. Importantly, the contact interface between Nb^traN^ and TraN^surf^ is rich in non-polar amino acids, suggesting their potential contribution to stable complex formation (**Fig. 3g**). To test this hypothesis, we introduced alanine substitutions into the interacting residues of CDR2 and CDR3 to disrupt the specific interaction, and then conducted macroscopic aggregation assays and isothermal titration calorimetry (ITC) to evaluate their impact on Nb^traN^/TraN binding. In line with our structural data, we did not observe any cell aggregates for either the CDR2 (F47A/Y58A) or CDR3 (Y98A/L100A) mutants when mixed with cells displaying TraN (**Supplementary Fig. 4a**). These mutants also displayed significantly reduced binding affinities for TraN (**Supplementary Fig. 4b**-e). Together, these data indicate that the hydrophobic residues within two of the CDRs of Nb^traN^ interact with the surface-exposed domain of TraN, facilitating cell-cell adhesion when Nb^traN^ is mounted on bacterial surfaces.

### Deep Mutational Nanobody Engineering Enhances the Binding Affinity of Nb^traN^ for <u>TraN</u>

Although we successfully identified a functional CAM targeting TraN from our nanobody pool, the binding affinity of Nb^traN^ was relatively modest (**Fig. 3d,e**). We postulate that nanobodies of higher affinity may not be present in the original library since the number of unique nanobody sequences was not saturated. This limitation in pool diversity could have impeded the discovery of high-affinity CAMs. Therefore, we set out to directly engineer Nb^traN^ by means of deep mutational scanning in conjunction with our whole-cell screening platform to obtain nanobody variants displaying an improved affinity for TraN (**Fig. 4a**). Our rationale was that if cells expressing the Nb^traN^ variants exhibited an enhanced affinity for TraN, they would acquire selective markers in a higher frequency, resulting in their enrichment within the population during the selection process. As an initial step, we focused on the nanobody’s CDR3 sequence, given that it is longer than other CDRs and is an important loop for antigen recognition^23^. Accordingly, we generated a focused library by single-site saturation mutagenesis of each residue in the CDR3 of Nb^traN^ using commercially available chip oligonucleotides (14 residues × 19 alternative amino acids/residues = 266 variants in the nanobody pool, as illustrated in **Fig. 4a**). The newly constructed nanobody library was then introduced into *E. coli* consisting of a display system and incubated with S17-1 cells expressing TraN for selection. After a single round of panning, we harvested surviving transconjugants on selective media and subjected them to high-throughput sequencing. We quantified the number of transconjugants by analyzing unique nanobody sequences in the pre- and post-selection populations.

**Fig. 4.**
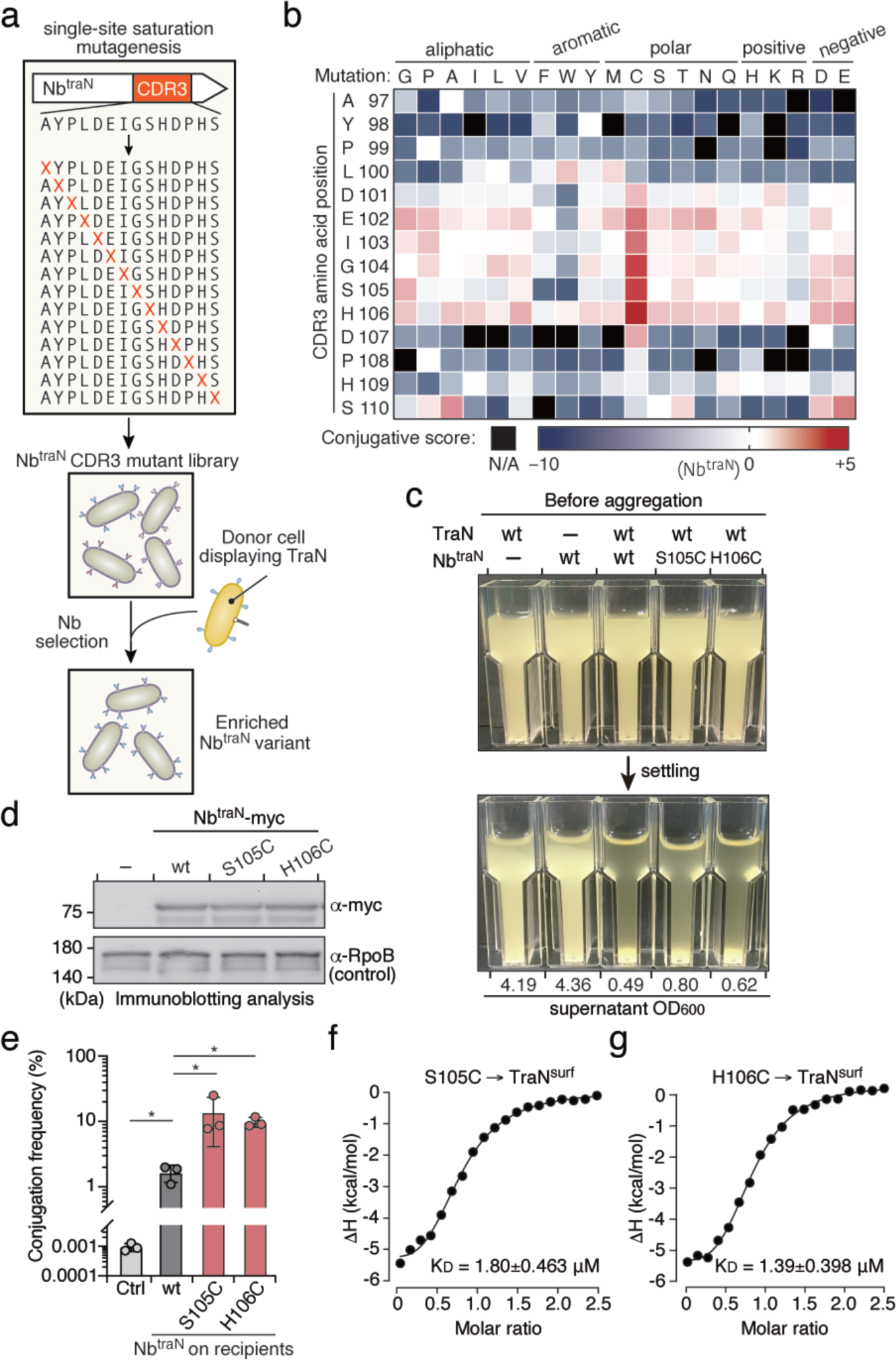
Discovery of optimized Nb^traN^ variants through deep mutational scanning. (**a**) Selection workflow of the deep mutational scanning. Plasmids carrying Nb^traN^ genes with single-site mutations in CDR3 were utilized for library construction. The mutant library underwent a single round of selection with donor cells displaying TraN to enrich for bacteria expressing potent Nb^traN^ variants displaying stronger binding capacity. (**b**) The conjugative scores of individual Nb^traN^ CDR3 mutations are depicted in a heatmap. The X-axis represents individual amino acid mutation, and the Y-axis represents the residue position of CDR3 in wild-type Nb^traN^. Mutations not identified after selection were excluded from the analysis and are shown in black (N/A). (**c**) Macroscopic aggregation analysis to assess cell-cell adhesion between *E. coli* strains displaying TraN and the indicated Nb^traN^ variants. The OD_600_ of the culture supernatant is shown below the image. The cells expressing TraN or wild-type Nb^traN^ were used as controls. (**d**) Immunoblotting analysis of *E. coli* expressing the Nb^traN^ variants carrying a C-terminal myc tag after a 16-hour incubation. The loading control is *E. coli* RNA Polymerase β protein (RpoB). Migrations of a size standard, in kDa, are indicted (left). (**e**) Conjugation frequency of pGenR under the liquid growth condition. Recipient cells expressing indicated nanobodies were co-cultured with donor cells expressing TraN. Data are presented as means ± SD. n = 3 technical replicates, representative of two biological replicates. *p < 0.05, t-test. (**f, g**) The ITC binding profiles of the indicated Nb^traN^ variants with TraN^surf^ at 25 °C. See also **Supplementary Fig. 5** and 6.

To gain a comprehensive understanding of the amino acid variants within the CDR3 of Nb^traN^ that affect conjugation in our screening process, we generated a sequence-function map illustrating variant conjugative scores (**Fig. 4b and Supplementary Fig. 5a**). Compared to wild-type Nb^traN^, 69% of the variants showed a decrease in conjugation ability following screening, whereas 31% showed an increase. Notably, substitutions of residue Y98 in CDR3 with other amino acids resulted in a reduced number of transconjugants in the pool, aligning with our above-described findings regarding its role in mediating the interaction between Nb^traN^ and TraN (**Supplementary Fig. 4**). Several Nb^traN^ variants significantly outperformed wild-type Nb^traN^ in the whole-cell screening platform. We were particularly interested in the cysteine variants located sequentially from residues 101 to 107, all of which elicited a substantial increase in the transconjugant population. These residues are positioned in close proximity to the Nb^traN^/TraN^surf^ interface in our complex structure (PDB: 8X7N) and have the potential to interact more effectively with TraN upon mutating to cysteine during cell-cell contacts (**Supplementary Fig. 5b**). Again, using macroscopic cell aggregation assays, we confirmed that cells displaying the CDR3 cysteine variants with the highest conjugative score (S105C and H106C) presented precipitation upon mixing with TraN-expressing cells (**Fig. 4c and Supplementary Fig. 6a**). To rule out indirect effects related to differences in protein levels, we conducted immunoblotting analysis and did not observe significant increases in protein expression for the cysteine variants relative to wild-type Nb^traN^ (**Fig. 4d**). We further assessed if cells displaying the Nb^traN^ cysteine variants could promote cell-cell adhesion through comparative conjugation in liquid culture. Consistent with the results from our deep-mutational screening, recipients displaying the Nb^traN^ cysteine variants exhibited a more than 5-fold increase in conjugation frequencies when incubated with S17-1 donor cells expressing TraN (**Fig. 4e**). Finally, we purified Nb^traN^S105C and Nb^traN^H106C and used ITC to measure their binding affinities for TraN^surf^. Compared to wild-type Nb^traN^, the cysteine variants presented, on average, a 3-fold increase in binding to TraN^surf^ *in vitro* (**Fig. 4f,g and Supplementary Fig. 6b**,c). In summary, we have demonstrated that our whole-cell screening platform not only identifies functional CAMs from a synthetic library, but this method is feasible for enhancing the cell adhesion capabilities of those identified CAMs by deploying strategic deep mutational engineering.

### Discovery of Nanobodies Targeting the Embedded Membrane Proteins OmpA and OmpC

Encouraged by the success of our selection method, we set out to identify functional CAMs capable of recognizing embedded membrane proteins. These proteins possess few surface-exposed epitopes and pose challenges for antibody acquisition through conventional screening methods or large-animal immunization (**Supplementary Fig. 7**). We selected two target antigens, OmpA and OmpC, representing two bacterial proteins naturally localized on the outer membrane that exert important roles in maintaining bacterial surface integrity and regulating membrane permeability^24, 25^. To our knowledge, no CAMs targeting the extracellular epitopes of OmpA or OmpC have been reported previously, making nanobodies targeting them especially unique and relevant. It is worth noting that although endogenous OmpA and OmpC are present in the S17-1 conjugative cells, we overexpressed these proteins by introducing an additional gene copy by plasmids, thereby enhancing the likelihood of isolating synthetic binders against these target antigens. Next, we incubated the S17-1 strains (OmpA^+^ or OmpC^+^) with the bacteria-displayed nanobody library, with the objective of identifying functional CAMs specifically tailored to these embedded membrane proteins.

Following three rounds of bio-panning along with sequencing analyses, we successfully identified nanobodies targeting both of these membrane proteins, which we have designated Nb^ompA^ and Nb^ompC^ (**Fig. 5a**). We found that *E. coli* harboring OmpA and OmpC underwent self-aggregation when they expressed these nanobodies on their surfaces, indicating their potential utility as CAMs (**Fig. 5b**). A comparative conjugation assay in liquid culture supported this notion, as the recipient cells displaying either Nb^ompA^ or Nb^ompC^ on their surfaces presented greatly enhanced conjugation frequencies upon incubation with S17-1 donor cells having OmpA and OmpC antigens (**Fig. 5c,d**). Finally, to validate the binding capacity of Nb^ompA^ and Nb^ompC^ to their corresponding surface antigens, we explored their potential utility as reagents for immunofluorescence labeling. We fused these nanobodies in-frame with a green fluorescent protein, referred to as Nb-GFP, purified them, and then performed a standard cell labeling procedure for visualization by fluorescence microscopy (**Fig. 5e,f and Supplementary Fig. 8**). We observed distinct fluorescence signals colocalized with wild-type *E. coli* hosting endogenous OmpA and OmpC when bound with Nb^ompA^-GFP and Nb^ompC^-GFP, respectively. These signals diminished when the Nb-GFPs were incubated with mutant *E. coli* strains (Δ*ompA*, Δ*ompC*). These results not only evidence the binding capacity of the Nb-GFPs toward their cognate membrane antigens, but also highlight the potential utility of CAMs for detecting bacteria expressing desired surface proteins.

**Fig. 5.**
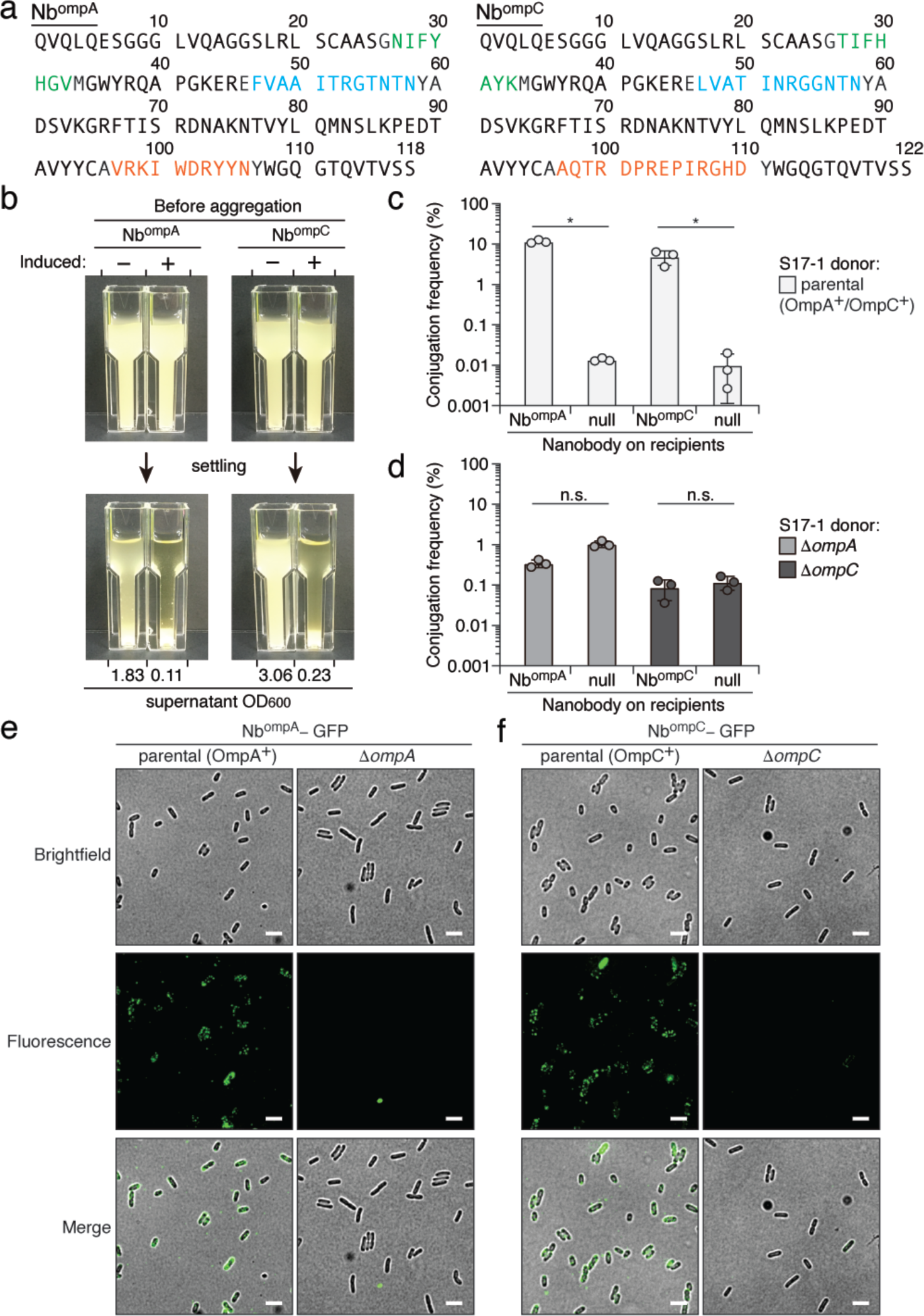
Nanobody CAM targeting of *E. coli* integral membrane proteins OmpA and OmpC. (**a**) The amino acid sequence of Nb^ompA^ and Nb^ompC^. CDR1 (green), CDR2 (red), and CDR3 (blue) are indicated. (**b**) Self-aggregation of *E. coli* expressing Nb^ompA^ and Nb^ompC^. Cultures underwent 24-hour aTc induction before settling. Cultures that did not undergo induction (*i.e.,* not expressing nanobodies) are presented as controls. (**c, d**) Conjugation frequency of pGenR under the liquid growth condition. Recipient cells expressing indicated nanobodies were co-cultured with S17-1 donor cells expressing (**c**) or not expressing (**d**) the indicated antigens. Data are presented as means ± SD. n = 3 technical replicates, representative of two biological replicates. *p < 0.05, t-test. (**e, f**) Fluorescence micrographs showing the antigen-binding specificity of Nb^ompA^-GFP (**e**) and Nb^ompC^-GFP (**f**). Nb^ompA^-GFP and Nb^ompC^-GFP were incubated with wild-type *E. coli* and the antigen-deleted strains (Δ*ompA* or Δ*ompC*) for 1 hour before observation. Scale bar = 1 μm. Full-size images are available in **Supplementary Fig. 8**.

### Nanobody-Programmed Inhibitor Cells Selectively Deplete Target Bacteria from Mixed Populations

One emerging and valuable application of synthetic CAMs is targeted antimicrobial interventions^8, 10^. We recently developed programmed inhibitor cells (PICs) displaying surface nanobodies that can selectively eliminate *E. coli* expressing cognate surface antigens within complex microbial communities, leveraging the potent antibacterial activity of the type VI secretion system (T6SS) in *Enterobacter cloacae*^26, 27^ (**Fig. 6a**). Despite being an important advance, the nanobodies employed in prior studies that recognize the embedded membrane protein BamA can be hindered by bacterial surface structures such as lipopolysaccharides (LPS)^8, 28^. Taking advantage of our newly identified CAMs, next we endeavored to determine if Nb^ompA^ and Nb^ompC^ are compatible with PICs for targeted cell elimination in bacterial mixtures. More importantly, we wanted to determine if PICs equipped with these nanobodies could selectively kill *E. coli* without altering native surface structures.

**Fig. 6.**
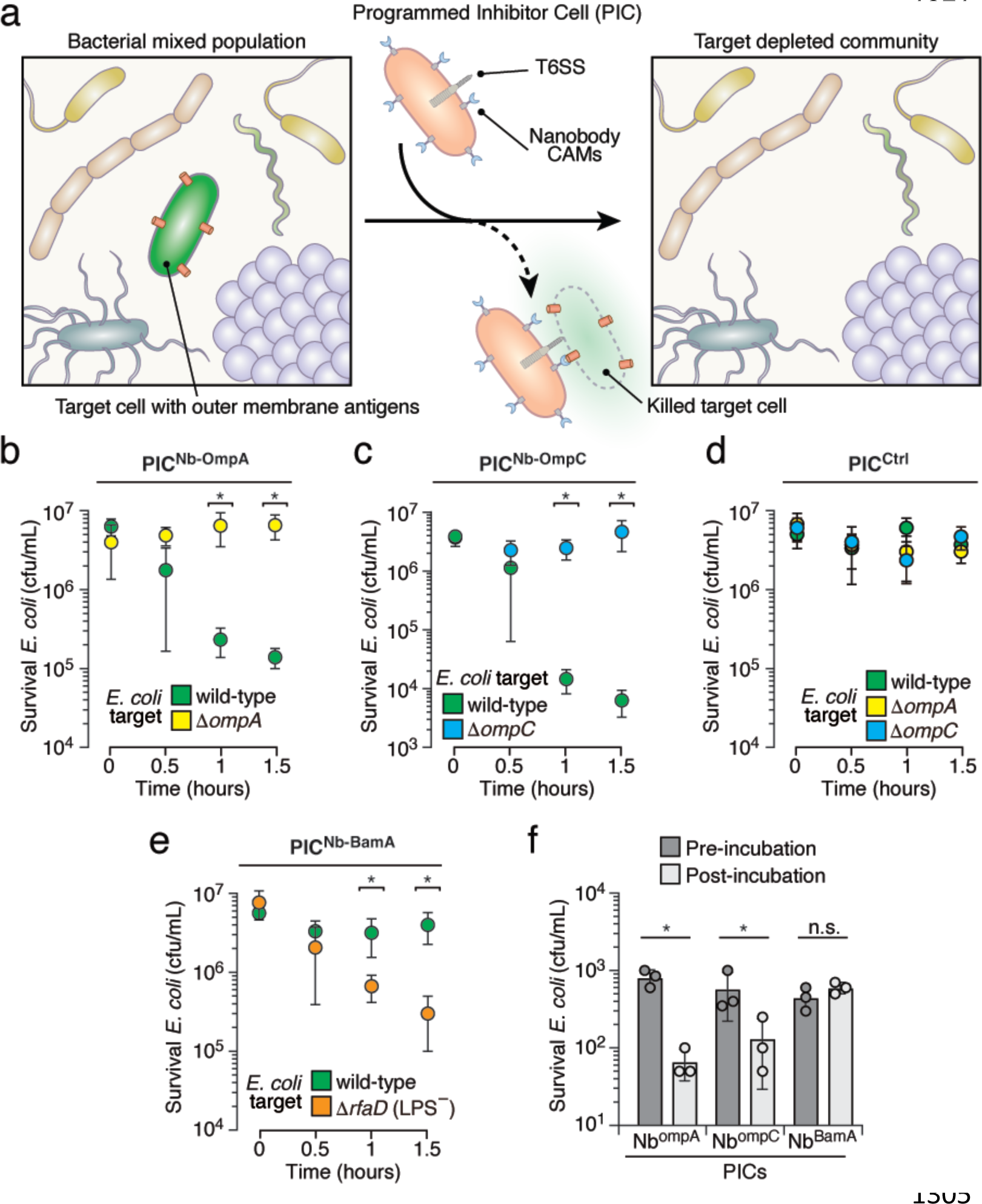
Mounting Nb^ompA^ and Nb^ompC^ on PICs depletes target *E. coli* from bacterial mixtures. (**a**) Schematic of target depletion of bacteria (green) from a complex bacterial population by PICs (pink). (**b-e**) Survival (CFUs/mL) of *E. coli* strains (wild-type, Δ*ompA*, Δ*ompC*, and Δ*rfaD*, colored green, yellow, blue, and orange, respectively) upon incubation with PICs displaying Nb^ompA^ (**b**), Nb^ompC^ (**c**), non-match control (Ctrl) (**d**), or Nb^BamA^ (**e**) in liquid medium (n = 3 technical replicates ± S.D.; *p < 0.05, t-test). Two biological repeats were conducted. (**f**) Survival (CFUs/mL) of wild-type *E. coli* within a mouse fecal bacterial population before and after incubation with PICs displaying Nb^ompA^, Nb^ompC^, or Nb^BamA^ (n = 3 technical replicates ± S.D., representative of two biological replicates. *p < 0.05, t-test). See also **Supplementary Fig. 9** and 10.

We proceeded to generate PICs displaying nanobodies that target OmpA and OmpC, denoted PIC^Nb-OmpA^ and PIC^Nb-OmpC^, respectively. To examine their capacity to direct the antibacterial activity of PICs against target *E. coli*, we performed target depletion assays using a high cell density and short timescale to mimic natural microbial communities found in the environment or host. Indeed, we found that both PIC^Nb-OmpA^ and PIC^Nb-OmpC^ effectively reduced the cell viability of *E. coli* within an hour by more than an order of magnitude (PIC^Nb-OmpA^, 27.7-fold; PIC^Nb-OmpC^, 168.2-fold), in a manner dependent on the presence of respective nanobodies, the T6SS, and the corresponding antigens on target cells (**Fig. 6b-d and Supplementary Fig. 9**). In contrast, PICs equipped with nanobody targeting BamA (PIC^Nb- BamA^) exerted a minimal impact on wild-type *E. coli* CFU numbers, unless LPS were truncated^8^ (**Fig. 6e and Supplementary Fig. 9**). These results provide evidence for the compatibility of both Nb^ompA^ and Nb^ompC^ with PICs to achieve targeted cell depletion. Moreover, we have showcased examples of specific antimicrobial interventions with minimal perturbation of the target cells. Lastly, to assess the feasibility of PIC^Nb-OmpA^- and PIC^Nb-OmpC^-mediated targeting within complex bacterial mixtures, we isolated total fecal bacteria from conventionally reared mice, which contain wild-type *E. coli* targets. We measured the capacity of PIC^Nb-OmpA^ and PIC^Nb-OmpC^ to selectively deplete *E. coli* from this diverse and undefined microbiome. Within 60 minutes post-PIC addition, *E. coli* CFU levels dropped significantly by 92% (PIC^Nb-OmpA^) or 78% (PIC^Nb-OmpC^), whereas PICs displaying Nb^BamA^ (PIC^Nb-BamA^) had no observable impact on survival of the target cells (**Fig. 6f and Supplementary Fig. 10**). This outcome underscores the potential of PIC-mediated targeting as a robust approach for selectively depleting target cells from complex microbial communities.

## DISCUSSION

Cell adhesion molecules are important in the study of fundamental biology and represent promising biotechnological tools. However, a key issue is how to identify functional CAMs that effectively mediate intercellular assemblies for desired outcomes. In this study, we report our development of a whole-cell screening platform designed to discover CAMs that facilitate programmable bacteria-bacteria adhesion. The success of this platform relies on a strategic combination of functional and positive selection, leveraging the T4SS for contact-dependent DNA delivery. The T4SS serves as a discriminatory enrichment tool, selectively favoring bacteria displaying functional CAMs, thereby addressing limitations associated with conventional methods. Our successful identification of nanobodies targeting outer membrane proteins attests to the effectiveness of our approach. The nanobodies isolated in this study exhibit stable and specific interactions with their cognate target antigens, underscoring their potential as versatile tools for engineering bacteria-bacteria adhesions. These CAMs present opportunities for future applications, such as engineering complex multi-component metabolic pathways and biomaterials^5-7^, as well as precision-targeted manipulation^8, 10^. Furthermore, we anticipate our method will prove compatible with existing cell-cell adhesion platforms, enabling the design and fine-tuning of multicellular assemblies, patterns, and morphologies, thereby positioning CAMs as key components in the synthetic microbiology toolkit^11, 12^.

Our study primarily serves to demonstrate the platform design and provide proof-of-concept applications, yet we recognize the potential for further optimization with a view toward specific targets. For example, we detected basal DNA conjugation frequencies in the absence of CAMs (0.01-0.001%, as shown in **Fig. 1 and 2**), indicating avenues for improvement. Our future efforts will concentrate on identifying or engineering alternative conjugative cells to maximize specific gene transfer efficiency while minimizing non-specific cell-cell adhesion. Importantly, the flexibility of our whole-cell screening platform, as exemplified through deep mutational engineering (**Fig. 4**), establishes a foundation for future endeavors aimed at expanding CAM diversity. Incorporating a more extensive and diverse pool of CAMs promises broader applicability and functionalities. Lastly, to extend the scope for identifying functional CAMs, future studies could explore alternative screening platforms utilizing Gram-positive bacteria, enabling the targeting of membrane antigens beyond the Gram-negatives examined herein. Despite divergences in DNA transfer mechanisms, the presence of conjugation machinery in Gram-positive bacteria supports the feasibility of extending contact-dependent DNA transfer via synthetic cell-cell adhesion^29^.

Despite the primary goal of this study being to develop a platform for isolating CAMs between bacteria, we envision that the principles of selection we deployed can transcend this specific scope. One notable advantage of our selection design is that by exploiting cell surface display systems to present antigens of interest. We have overcome technical challenges associated with recombinant protein purification in conventional screening platforms^14, 22^. Numerous bacterial surface display systems have been developed in recent decades, which can efficiently express heterologous proteins for various purposes^30^. We anticipate that our rapid whole-cell-based selection regime can be extended to isolate antigen-specific binders using those protein display systems.

Our study not only demonstrates how functional CAMs can be identified via a high-throughput screening workflow, but also showcases their compatibility with existing technologies relying on cell-cell adhesion. By installing isolated CAMs targeting OmpA or OmpC on the surface of PICs, we could specifically direct the antibacterial activity of PICs toward target bacteria in mixed populations (**Fig. 6**). Overall, our results herein highlight the vast potential of our screening platform and its potential integration with other synthetic biology tools. For instance, we envisage additional possibilities using CRISPR-based systems or engineered antimicrobials^10, 31-33^. Combining different synthetic biology approaches for the precise manipulation of microbial communities could yield synergistic benefits with diverse applications. The adaptability and versatility of CAMs identified using our platform offer opportunities for innovation, both in terms of navigating the challenges posed by biological system complexity and deciphering potential applications. In conclusion, this study sheds light on the transformative potential of CAMs in microbiology and synthetic biology.

## METHODS

### Bacteria, Yeast, and Culture Conditions

The strains utilized in this study are detailed in **Supplementary Table 2**. We used the following *Escherichia coli* strains: S17-1 ATCC 47055 to display antigens of interest and transfer plasmids containing distinct antibiotic-resistance markers for the selection method; MG1655 to display nanobodies and validate nanobody-binding specificity; DH5α and PIR1 in cloning and plasmid maintenance; S17-1 λ pir for conjugal transfer of pRE118 plasmid into *E. coli*; and BL21 (DE3) pRIL for overproduction and purification of protein samples. *Enterobacter cloacae* ATCC-13047 was utilized for target depletion assays^8^. All bacteria were routinely cultured at 37 °C in Luria Bertani (LB) medium unless noted otherwise. The yeast-display nanobody library was recovered by growing it in Yglc4.5 –Trp medium at 30 °C in a shaker^22^. Antibiotics and chemicals were used at the following concentrations: 50 μg/mL streptomycin; 150 μg/mL carbenicillin; 50 μg/mL kanamycin; 25 μg/mL chloramphenicol; 15 μg/mL gentamicin; 100 μg/mL trimethoprim; 500 ng/mL anhydrotetracycline (aTc); 100 μM isopropyl β-D-1-thiogalactopyranoside (IPTG); 1% (*w/v*) L-(+)-arabinose; and 200 mM sucrose.

### Mice

Fecal samples from three male wild-type C57BL/6J mice were utilized in this study. Care and maintenance of these mice were conducted according to the protocol of the Institutional Animal Care and Use Committee (IACUC) of the Institute of Molecular Biology (Academia Sinica, Taiwan) and approved by its Animal Center, where the mice were bred and housed. All mice were maintained under controlled light conditions (12-hour light-dark cycle) with access to standard food and water. Prior to oral gavage with wild-type *E. coli* MG1655 targets, the mice underwent a 4-hour fasting period.

### Plasmid Construction

The plasmids and primers utilized in our study are listed in **Supplementary Table 2**. Tet expression plasmids, which were used for nanobody display on the surface of both *E. coli* and *E. cloacae*, were derived from pDSG323-based expression plasmids as described previously^8, 11^. The TraN antigen expression plasmid (pDSG-TraN) was constructed by polymerase chain reaction (PCR) amplification and subsequent cloning of the *traN* gene (AAL23498.1) from the pSLT plasmid of *Salmonella* Typhimurium LT2 strain into pDSG323 at the *NotI* and *PstI* restriction sites. The OmpA and OmpC arabinose-inducible plasmids (pBAD18-OmpA, pBAD18-OmpC) were generated by PCR amplification and subsequent cloning of the respective genes (NP_415477.1, NP_416719.1) from the genomic DNA of *E. coli* MG1655 strain into the *SacI* and *XbaI* restriction sites of pBAD18. A Myc-tag sequence (EQKLISEEDL) was introduced at the C-terminus of Nb^traN^ and its variants by means of site-directed mutagenesis^34^.

For the TraN^surf^/Nb^traN^ complex purification constructs, the sequences were amplified and cloned into multiple cloning site (MCS)-1 (*BamH1* and *NotI* sites) and MCS-2 (*NdeI* and *XhoI* sites) of pETDuet-1, respectively, resulting in fusion of an N-terminal hexa-histidine to TraN^surf^. Purification of Nb^traN^, Nb^ompA^, and Nb^ompC^ involved amplifying their sequences and cloning them into the MCS (*NdeI* and *XhoI* sites) of pET22b, fused with a C-terminal hexa-histidine tag. Nb^traN^ nanobody variants were introduced into the expression plasmids using site-directed mutagenesis^34^. The Nb^ompA^-GFP and Nb^ompC^-GFP constructs were generated by PCR amplification of the GFP sequence from pET11a-GFP-avi and inserting it between the nanobody sequences and the hexa-histidine tag at the C-terminus of pET22b plasmids via Gibson assembly^35^. To purify TraN^surf^ alone, the sequence was amplified and cloned into MCS-1 (*BamH1* and *NotI* sites) of pETDuet-1, generating a fusion construct with an N-terminal hexa-histidine tag. The *E. coli ompW* in-frame deletion plasmid (pRE118_EcoliΔompW) was generated by utilizing the suicide vector pRE118^36^. We amplified 750 bp regions flanking the deletion, joined them by means of splicing by overlap extension (SOE) PCR, and subsequently cloned them into pRE118 at the *SacI* and *XbaI* restriction sites. For mobilizable plasmids with distinct antibiotic-resistance genes, the pGenR was created from plasmid pPSV37 by deleting the T7-expression system and re-ligating the vector by means of Gibson assembly^37^, whereas pTmpR was created from plasmid pSCrhaB2 by deleting the rhamnose-expression system^38^. The pUC18T-mini-Tn7T plasmid consists of an ampicillin-resistance gene and its oriT was renamed as pAmpR for the purposes of our study^39^.

### Genetic Manipulation

We used the lambda red recombinase system to generate *E. coli* strains in which *ompA*, *ompC*, or *trbE* had been deleted^40^. In brief, PCR products comprising a kanamycin resistance cassette from pKD4, flanked by 50∼100 bp homologous to the 5’ and 3’ termini of these genes, were electroporated into *E. coli* strains (MG1655 or S17-1) carrying pKD46 and induced by arabinose for 5 hours at 30 °C to express the recombinase. Subsequently, *E. coli* was incubated in LB for 1 hour, plated on chloramphenicol-containing LB agar, and incubated overnight at 37 °C. To remove the kanamycin resistance cassette from *E. coli*, pCP20 plasmid hosting the FLP recombinase gene was transformed into the deletion strains, which were then grown at 30 °C on carbenicillin-containing LB agar. A single colony was picked and inoculated into LB media at 43 °C to select for loss of pCP20.

To delete *ompW* from *E. coli* MG1655, pRE118_EcoliΔompW was transformed into *E. coli* S17-1 λ pir. The donor cells carrying the deletion construct and *E. coli* MG1655 were grown overnight on LB plates containing antibiotics, as appropriate. The cells were then scraped together to create a 2:1 donor-recipient mixture, which was spread out on an LB agar plate and incubated at 37 °C for 6 hours to facilitate plasmid transfer via conjugation. Cell mixtures were then scraped into LB and plated onto LB medium agar plates supplemented with kanamycin and chloramphenicol to select for *E. coli* containing the chromosomally inserted deletion construct. The *E. coli* merodiploid strains were grown overnight in non-selective LB medium at 37 °C, followed by counter-selection on LB low salt plates containing sucrose. Kanamycin-sensitive colonies were screened for allelic replacement by colony PCR and mutations were confirmed by Sanger sequencing of the PCR products.

### Bacterial Conjugation Assays

For conjugation assays, the conjugative strain *E. coli* S17-1 ATCC 47055 was employed as the donor carrying transferable plasmids with selection markers, whereas *E. coli* MG1655 served as the recipient. Overnight bacterial cultures were centrifuged at 21,100 x *g* for 1 minute to pellet the cells, followed by the removal of the culture supernatant. The cells then underwent a single wash in LB broth and were subsequently resuspended in LB broth. The donor and recipient cells were adjusted to OD_600_ = 0.005 and 0.0025, respectively. Liquid conjugation assays involved incubating cell mixtures at 37 °C with continuous agitation at 200 rpm for 6 hours. For conjugation assays on solid media, cell mixtures were spotted onto LB agar plates and incubated at 37 °C for 6 hours. Following incubation, the cells were collected, serially diluted, and plated on selective media to quantify colony-forming units (CFUs). Three transferable plasmids contained gentamicin-, ampicillin-, or trimethoprim-resistance genes, respectively. The recipient strain was marked by chromosomal insertion of a chloramphenicol-resistance gene, whereas the donor strains remained unmarked. The total recipient population was assessed by enumerating CFUs on LB plates containing chloramphenicol. The count of transconjugant populations was determined by enumerating CFUs on LB plates with chloramphenicol, which also contained the relevant resistance markers (gentamicin, ampicillin, or trimethoprim), depending on the specific conjugative plasmid employed in the selection process.

### Ag/Nb Mismatch Conjugation Experiment

We used four different mixed cell cultures in the mismatch conjugation assay. Conjugative *E. coli* S17-1 strains displaying either Ag1 or AgInt from Tet expression plasmids were co-cultured with recipient strains displaying Nb1 or NbIB10 from Tet expression plasmids. Initial cell concentrations for donor and recipient strains were set to OD_600_ = 0.005 and 0.0025, respectively. Following 6-hour incubation in a liquid medium containing kanamycin and aTc with continuous shaking at 37 °C, the cells were subjected to serial dilution and plated on selective media to quantify CFUs as described above.

To assess if the whole-cell screening approach exhibits specificity toward low-abundance target cells within mixed bacterial populations, overnight cell cultures were centrifuged at 21,100 x *g* for 1 minute. The culture supernatant was discarded, and the cell pellet was collected. After a single wash, cells were resuspended in LB broth, and their concentration was adjusted to OD_600_ = 1.0. Recipient cells expressing Nb1 and Null were mixed at varying ratios (10^1^- to 10^7^-fold dilutions as specified). The recipient mixture and donor cells displaying Ag1 were diluted to OD_600_ = 0.0025 and 0.005, respectively. The assay comprised three rounds of bio-panning. In the initial round, the donor strain carrying a transferable gentamicin resistance gene was utilized. Following 6-hour incubation at 37 °C with shaking at 200 rpm, cultures were diluted and plated on selective media for CFU quantification. The remaining culture was plated on selective media to initiate the next panning round. After 16 hours of incubation at 37 °C, recipient colonies that grew on selective plates were harvested and resuspended in LB broth for the second panning round. The recipient culture and the donor carrying a transferable ampicillin resistance gene were diluted to OD_600_ = 0.0025 and 0.005, respectively. After incubation again at 37 °C with agitation at 200 rpm for 6 hours, cultures were diluted and plated on selective media for CFU quantification. The remainder of the culture was plated on selective media for the subsequent panning round. Following another 16 hours of incubation at 37 °C, recipient colonies that grew on the selective plates were harvested and resuspended in LB broth for the third panning round. The recipient culture and the donor carrying a transferable trimethoprim resistance gene were diluted to OD_600_ = 0.0025 and 0.005, respectively. After incubation at 37 °C with agitation at 200 rpm for 6 hours, cultures were diluted and plated on selective media for CFU quantification.

### Bacterial Nanobody Display Library Construction

The bacterial display library used in this study was derived from a yeast nanobody library established in a previous study^22^. Plasmids containing nanobody genes were first extracted from the yeast library, and the nanobody sequences were subsequently amplified via PCR. These fragments were then subcloned into pDSG323 backbone vectors using the Gibson assembly method and chemically transformed into *E. coli* DH5α for plasmid enrichment. A total of fifty separate Gibson assembly reaction products were transformed, resulting in the collection of >10^7^ colonies, after which plasmids were extracted and pooled. The bacteria nanobody display library (100 ng) was then transformed into the *E. coli* MG1655 *ΔompW* strain via electroporation. We selected the Δ*ompW* strain to minimize the interactions between OmpW and TraN, as suggested by Low *et al.*^21, 41^. The cells were harvested and resuspended in LB with 10% glycerol for storage at -80 °C.

To construct the Nb^traN^ CDR3 library for deep mutational engineering, 14 sites on CDR3 sequences (AYPLDEIGSHDPHS) underwent mutagenesis, with each site mutated into all desired amino acids using optimal *E. coli* codons. GenScript performed DNA oligonucleotide library synthesis on a programmable column. The pooled mutagenesis library was subsequently cloned into the pDSG323 backbone and transformed into *E. coli* DH5α competent cells. Sanger sequencing was conducted to determine library capacity before colonies were collected, and the plasmids were then extracted. The pool of plasmids was then transformed into the *E. coli* MG1655 Δ*ompW* strain via electroporation, resulting in a collection of >30,000 colonies stored in LB broth with 10% glycerol at -80 °C.

### Nanobody CAM Isolation using the Whole-Cell Screening Platform

Conjugative *E. coli* S17-1 expressing indicated antigens (TraN, OmpA, or OmpC) along with transferring plasmids containing distinct antibiotic-resistance markers served as the donor strain, whereas the bacterial nanobody library acted as the recipient. In the initial bio-panning round, the donor with a mobilized gentamycin-resistance plasmid and the recipient nanobody display library were both diluted to OD_600_ = 0.05 in LB broth and incubated at 37 °C with shaking at 200 rpm until log phase was attained. The donor cells and the library culture were then diluted to OD_600_ = 0.0025 and 0.005, respectively, and co-cultured in LB broth with aTc. After 6-hour incubation at 37 °C with shaking at 200 rpm, the culture was plated on selective media containing gentamycin and chloramphenicol and incubated at 37 °C for 16 hours. All colonies (>10^6^ clones) were harvested and resuspended in LB broth. In the second round of panning, recipient cells from the first round and donor cells carrying a transferrable ampicillin resistance plasmid were mixed at OD_600_ = 0.0025 and 0.005, respectively. Following the same panning conditions as in the initial round, the culture was plated on selective media containing carbenicillin and chloramphenicol. After 16-hour incubation at 37 °C, all colonies were collected and resuspended in LB broth for the third round of panning. The culture harvested after the second panning round was co-cultured with donor cells carrying a transferrable trimethoprim resistance plasmid at a ratio of OD_600_ = 0.0025 and 0.005, respectively. After 6-hour co-culture under the same conditions as the previous round, the culture was plated on selective media containing trimethoprim and chloramphenicol. All grown colonies were harvested. Cells obtained after each panning round were collected to analyze nanobody diversity at each time point. For Nb^traN^ deep mutational engineering, conjugative *E. coli* S17-1 expressing TraN served as the donor strain, and the bacterial Nb^traN^ mutant library served as the recipient. The selection process comprised a single round of bio-panning, as described above.

### Macroscopic Cell Aggregation Assays

*E. coli* displaying nanobodies or antigens were initially inoculated in LB broth with aTc inducer. Following a 24-hour incubation at 37 °C, the cultures were vortexed and mixed in a 1:1 ratio with the indicated testing combination. The cell mixtures were then placed in cuvettes with a 1-cm light path at room temperature. After 24 hours of settling, cultures from the top 25% of the wells were collected, and their OD_600_ was measured using a Nano300 microspectrophotometer.

### Protein Purification

His-tagged TraN^surf^ and Nb^traN^ proteins were co-expressed from pET-Duet-1 in *E. coli* BL21 (DE3) pRIL. Stationary phase cultures of expression strains were used to inoculate 1 L of LB broth containing carbenicillin and chloramphenicol. Cells were cultured in a shaking incubator at 37 °C and 220 rpm until they reached log phase (OD_600_ = 0.6∼0.8), induced by IPTG, and then incubated at 30 °C and 220 rpm for 18 hours. Cells were harvested by centrifugation at 4 °C and 4,000 x *g* for 20 minutes and then resuspended in 40 mL lysis buffer (20 mM Tris-Cl pH = 7.5, 300 mM NaCl, 10 mM imidazole pH = 7.0, 1 mM DTT, protease inhibitor cocktail). Resuspended cells were disrupted with a microfluidizer and the cell debris was removed at 20,000 x *g* for 30 minutes at 4 °C. The Nb^traN^/TraN^surf^ protein complex was then purified from the supernatant individually by gravity flow through a 2 mL Ni-NTA agarose column. Bound proteins were then eluted using a linear imidazole gradient to a final concentration of 300 mM. The purity of each protein sample was assessed by 15% SDS-PAGE followed by Coomassie Brilliant Blue staining. Protein samples were further purified by fast protein liquid chromatography (FPLC) using gel filtration on a HiLoad 16/600 Superdex 200pg column (**Supplementary Fig. 3b**).

For protein binding experiments requiring free TraN^surf^ and nanobodies, proteins were overproduced in *E. coli* strain BL21 (DE3) pRIL transformed with the IPTG expression vectors containing the desired genes. Cultures were grown in LB medium at 37 °C to log phase (OD_600_ = 0.6∼0.8) and induced by IPTG. After induction at 30 °C, the cells were precipitated and resuspended in lysis buffer as described above. Resuspended cells were disrupted using a microfluidizer. The cellular debris was removed by centrifugation at 20,000 x *g* for 30 minutes at 4 °C. The histidine-tagged proteins were purified from the supernatant by gravity flow through a 2 ml Ni-NTA agarose column. Bound proteins were then eluted using a linear imidazole gradient to a final concentration of 300 mM. All protein samples were further purified by FPLC using gel filtration on a HiLoad 16/600 Superdex 200pg column. Fractions with high purity were concentrated and used in binding assays.

### Crystallization and Structure Determination

For crystallization of the Nb^traN^/TraN^surf^ complex, the Ni-NTA-purified protein was dialyzed in low salt buffer (5 mM Tris-HCl pH = 7.0, 150 mM NaCl, 0.5 mM TCEP) and concentrated to ∼20 mg/mL by spin filtration (10 kDa cutoff, Millipore). The concentrated sample was subsequently screened against commercially available crystallization screens from Hampton Research. Diffraction-quality crystals were successfully generated at 20 °C via hanging drop vapor diffusion in three weeks, maintaining a buffer-to-protein ratio of 1:2. The crystallization solution, conducive to crystal formation, consisted of 1 M ammonium sulfate and 0.1 M sodium acetate at pH = 4.6.

Cryo-protection of Nb^traN^/TraN^surf^ complex crystals was performed in 20% (*v/v*) glycerol in the crystallization solution described above. The X-ray diffraction datasets were collected on beamline TPS-07A at the National Synchrotron Radiation Research Center (Taiwan). X-ray diffraction intensities were processed using HKL 2000^42^. Phase determination of the Nb^traN^/TraN^surf^ complex was carried out using molecular replacement by Phaser in PHENIX, with poly-alanine nanobody (PDB: 5VNV) as a search model^43^. The Nb^traN^/TraN^surf^ complex model was built into the electron density map and further refined by Coot and PHENIX, respectively^43, 44^. Crystals were cubic (space group P212121), with five complexes per asymmetric unit. There are no residues in the unfavored region of the Ramachandran plots. Data collection and refinement statistics are shown in **Supplementary Table 1**.

### Fluorescence Microscopy

Bacterial cultures were grown overnight in M9 minimal medium at 37 °C with 200 rpm shaking. The cells were harvested by centrifugation at 8,000 x *g* for 1 minute. The culture supernatant was removed and washed with Phosphate-buffered saline (PBS) by spinning at 8,000 x *g* for 1 minute. The washed cells were resuspended in PBS containing 1% paraformaldehyde (PFA) for 1-hour cell fixation at room temperature. The fixed cells were subjected to three PBS washes. The cell suspensions were then adjusted to OD_600_ = 1.0 using PBS. The addition of 2 µM of Nb-GFPs to the cell suspensions preceded a 1-hour incubation at 37 °C with shaking at 200 rpm for nanobody binding. The cells were then washed four times with PBS at 6,000 x *g* for 1 minute. Cell suspensions were spotted on a 1.5% agarose pad settled on a microscope slide. Images were captured using a DeltaVision Core Deconvolution Microscopy system with an Olympus IX71 microscope and a 60x 1.42 NA oil-immersion objective.

### Determination of Nanobody Library Genetic Diversity

To analyze the diversity of the nanobody population within each round of the selection process, cells were harvested and total plasmids were extracted for targeted amplicon sequencing^45^. Nanobody genes were amplified through 20 cycles of PCR using specific primers (**Supplementary Table 2**). Eight unique molecular identifiers (UMI) were incorporated before the 5’ primers to eliminate duplicate molecules resulting from PCR amplification. The PCR amplicons with a gel size of around 400 bp were selected and extracted using a QIAquick Gel Extraction Kit. The purified amplicons were then quantified using Qubit DNA HS reagent, and the 400 bp size amplicons were verified using a Bioanalyzer 2100 system (Agilent Technologies).

Next, Index 1 (i7) and Index 2 (i5) sequences were added to the cleanup samples to generate uniquely tagged libraries by employing the Illumina DNA/RNA UD Indexes Set B, followed by eight cycles of PCR using the Index primers and KAPA HiFi HotStart Ready Mix. The indexed amplicons were subsequently cleaned up by Ampure XP reagent and quantified. The final library was quantified to confirm the expected size of around 489 bp, and the concentration was calculated using an Illumina pooling calculator. The library samples were then diluted to 2 nM and submitted for Illumina MiSeq sequencing, which was performed by the Genomics Core of the Institute of Molecular Biology (Academia Sinica, Taiwan).

Raw data analysis was conducted as follows. Initially, nanobody sequences were identified through the 5’ and 3’ primers, as detailed in **Supplementary Table 2**. Reads with the same UMI sequences were considered to have originated from the same original molecule, enabling the removal of duplicate reads that arise during library amplification steps and ensuring more accurate quantification of the diversity and abundance of unique molecules in the samples. The variable region was subsequently determined by scrutinizing the 25-bp segments upstream and downstream of conserved sequences. Alignment of each conserved sequence with the nanobody was achieved using the Smith-Waterman algorithm for local sequence alignment^46^. The variable region was delineated based on the alignments of conserved sequences at both ends. In cases where the number of mismatches exceeded five, the variable region was deemed absent. Lastly, all variable regions underwent clustering, and their frequencies were tallied (**Supplementary Table 3**).

### Conjugative Scores Measurement

To analyze the deep mutational scanning data, log2 enrichment ratios of mutations (*Mut*) were normalized by subtracting the enrichment ratios of the wild-type (*Wt*) sequence according to the following formula (**Supplementary Fig. 5a and Supplementary Table 3**):

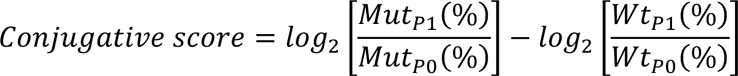

### Target Cell Depletion Assay

The target cell depletion assays were carried out as described previously^8^. In brief, overnight PIC cultures were diluted to OD_600_ = 0.05 and grown at 37 °C for 4 hours in LB broth with aTc to induce surface-exposed nanobodies (Nb^ompA^, Nb^ompC^, or Nb^BamA^). The cells were then pelleted, washed, and resuspended in LB, and then normalized to OD_600_ = 2. Mixtures of PICs and the indicated *E. coli* strains were then established at a 100:1 *v/v* ratio in LB broth with aTc. Bacterial mixtures were incubated at 37 °C with shaking at 200 rpm, and samples were collected at 0, 30, 60, and 90 minutes, followed by serial dilution and plating on selective media to quantify CFUs. PICs were distinguished by plating on LB containing streptomycin (intrinsic resistance), whereas *E. coli* were plated on LB containing chloramphenicol (chromosomal insertion of the chloramphenicol acetyltransferase gene).

For target depletion experiments in undefined bacterial communities, three reared C57BL/6J mice were fed by 200 μL oral gavage in 20% (*w/v*) sucrose solution containing 10^9^ CFUs of log-phase wild-type *E. coli* MG1655. Following ingestion of the bacterial suspension, the mice were provided with both standard rodent food and sterilized water. Fecal samples were collected at 24 hours post-feeding and suspended in 6 mL of PBS, and then homogenized using a tissue-tearor with a probe diameter of 7 mm (BioSpec). The samples were then gently added to the top of 2 mL of 80% (*w/v*) Nycodenz, followed by ultra-centrifugation at 10,000 x *g* for 40 minutes. The top layer of PBS solution was carefully removed, and the high-density fecal bacterial community from the middle layer was collected and normalized to OD_600_ = 3 in PBS. The resulting fecal bacterial community was mixed with an equal volume of PICs (OD_600_ = 2.0) expressing Nb^ompA^, Nb^ompC^, or Nb^BamA^ as described above. The bacterial mixtures were incubated at 37 °C with shaking at 200 rpm for 1 hour. Pre- and post-incubation samples were plated on LB with chloramphenicol or streptomycin to selectively quantify *E. coli* or PIC populations, respectively.

### Analysis of Protein Expression Levels

To analyze the expression level of nanobodies, *E. coli* expressing Nb^traN^ (wild-type and variants) were grown in LB medium supplemented with or without aTc at 37 °C for 6 hours and harvested at an OD_600_ of 1.0. For each quantification assay, cell pellets were resuspended in SDS-PAGE sample loading buffer. Samples were boiled at 100 °C for 10 minutes and equal volumes were loaded for SDS-PAGE, before being transferred to nitrocellulose membranes. Membranes were blocked in TBST (10 mM Tris-HCl pH 7.5, 150 mM NaCl, 0.1% (*w/v*) Tween-20) with 5% (*w/v*) non-fat milk for 30 minutes at room temperature, followed by incubation for 1 hour at room temperature with primary antibodies (anti-c-myc or anti-RNA Polymerase β) diluted 1:1000 in TBST. Blots were then washed by TBST, followed by incubation for 30 minutes at room temperature with the secondary antibody (anti-mouse IgG HRP conjugated) diluted 1:5000 in TBST. Finally, blots were washed again by TBST, developed, and visualized using a UVP Biospectrum 815 system.

### Isothermal Titration Calorimetry (ITC)

Titration experiments were performed using a Microcal PEAQ-ITC automated system (Malvern) at 25 °C. The ITC sample cell contained 390μL of 40μM TraN^surf^ in Tris buffer (20 mM Tris-Cl pH = 7.5, 150 mM NaCl, 0.5 mM TCEP). A syringe containing 125 μL of 500 μM Nb^traN^ or nanobody variants (Nb^traN^_F47A/Y58A, Nb^traN^_Y98A/L100A, Nb^traN^_S105C, and Nb^traN^_H106C) in the same Tris buffer was used to inject into the sample cell using 20x injections. The raw experimental data were analyzed using the MicroCal PEAQ-ITC Analysis software package provided with the instrument. The fitting function in the software was generated by clicking the iteration button until an excellent fit to the experimental data points of the isotherms had been obtained. The final parameters, including K_D_, changes in enthalpy (ΔH), Gibb’s free energy (ΔG), and entropy (ΔS), were generated by the same software.

### Quantification and Statistical Analyses

Statistical significance in bacterial competition experiments was assessed by unpaired t-tests between relevant samples. Details of statistical significance are provided in the figure legends.

## ACKNOWLEDGMENTS

We thank Drs. Marcos de Moraes, Yu-Chan Chao, and Ting laboratory members for helpful discussions and critical reading of the manuscript; Drs. Joseph Moguous, Yun Mou, Shu-Jung Chang, Hsin-Hung Chou, Liuh-Yow Chen, Jen-Hsuan Wei, Jun-Yi Leu, and Sue Lin-Chao for sharing reagents; Dr. Soo-Chen Cheng for sharing equipment; Drs. Shu-Yun Tung, Hsin-Nan Lin, Chen-Hsin Yu, Ching-Hung Tseng, and Xin-Jie Huang for technical assistance; and Dr. John O’Brien for manuscript editing. We acknowledge funding support from Academia Sinica (AS-CDA-112-L05 to S.-Y.T., AS-IVA-112-L05 to K.-C.H.) and the National Science and Technology Council, Taiwan (111-2311-B-001-006-MY2 and 112-2311-B-001-020 to S.-Y.T., 112-2320-B001-005-MY2 to W.-L.W., 112-2320-B-001-006-MY3 to K.-C.H.).

## SUPPLEMENTAL FIGURE LEGENDS

**Fig. S1.**
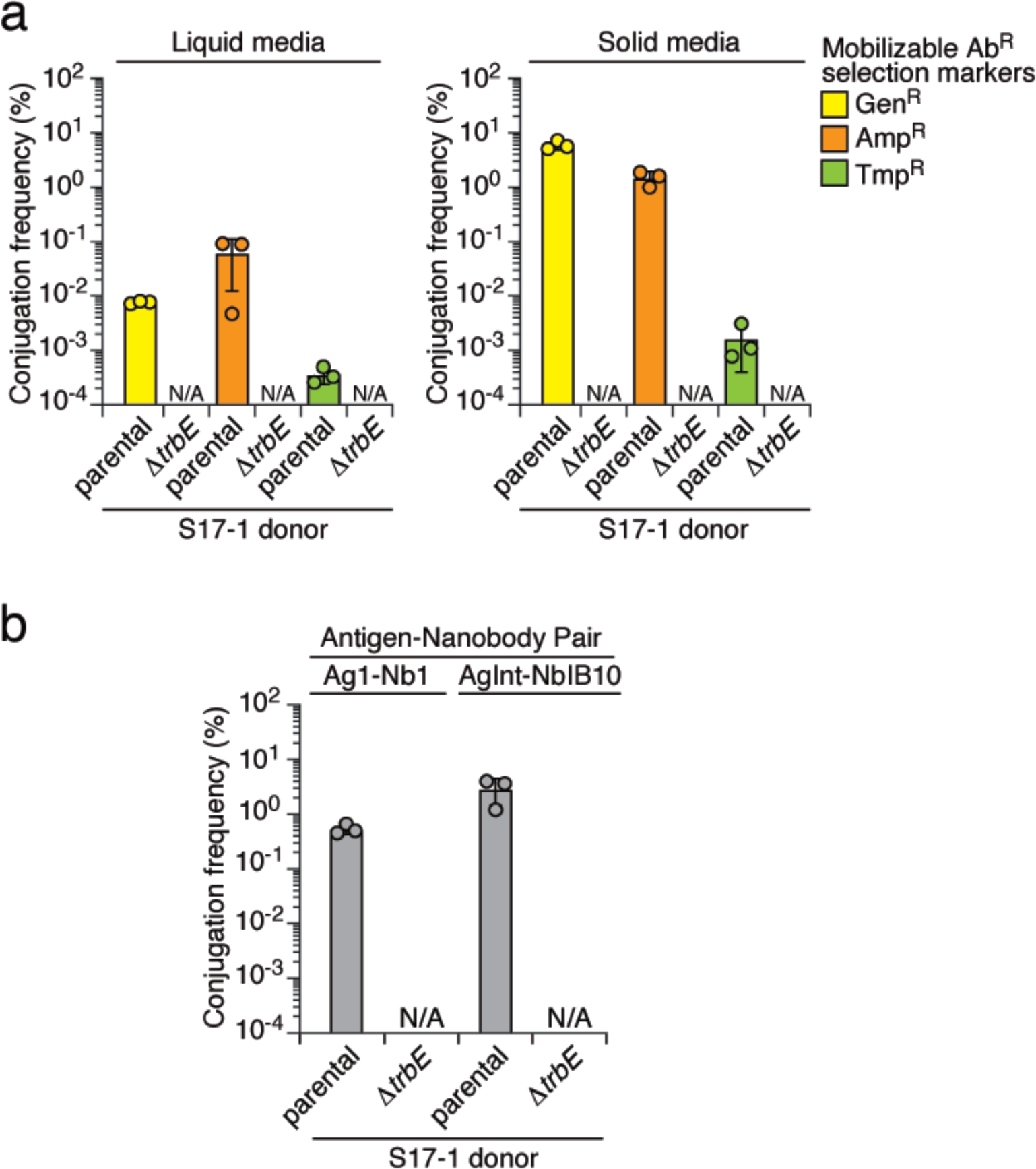
Dependency of selection marker gene transfer on a functional T4SS. (**a**) The conjugation frequency of selection markers from the WT and Δ*trbE* (dysfunctional T4SS) donors to recipients. Conjugation assays were performed under both liquid and solid growth conditions, utilizing pGenR, pCarbR, and pTmpR as selection markers. (**b**) The impact of a deficient T4SS on the donor (*E. coli* S17-1) to convey the selection marker (pGenR) to the recipient (*E. coli* MG1655). Two binding pairs were employed: Ag1 and AgInt were displayed by the donors, and their corresponding nanobodies (Nb1 and NbIB10) were displayed by the recipients. Data are presented as means ± SD. n = 3 technical replicates, representative of two biological replicates. *p < 0.05, t-test. Conjugation frequency lower than the detection limit designated as N/A.

**Fig. S2.**
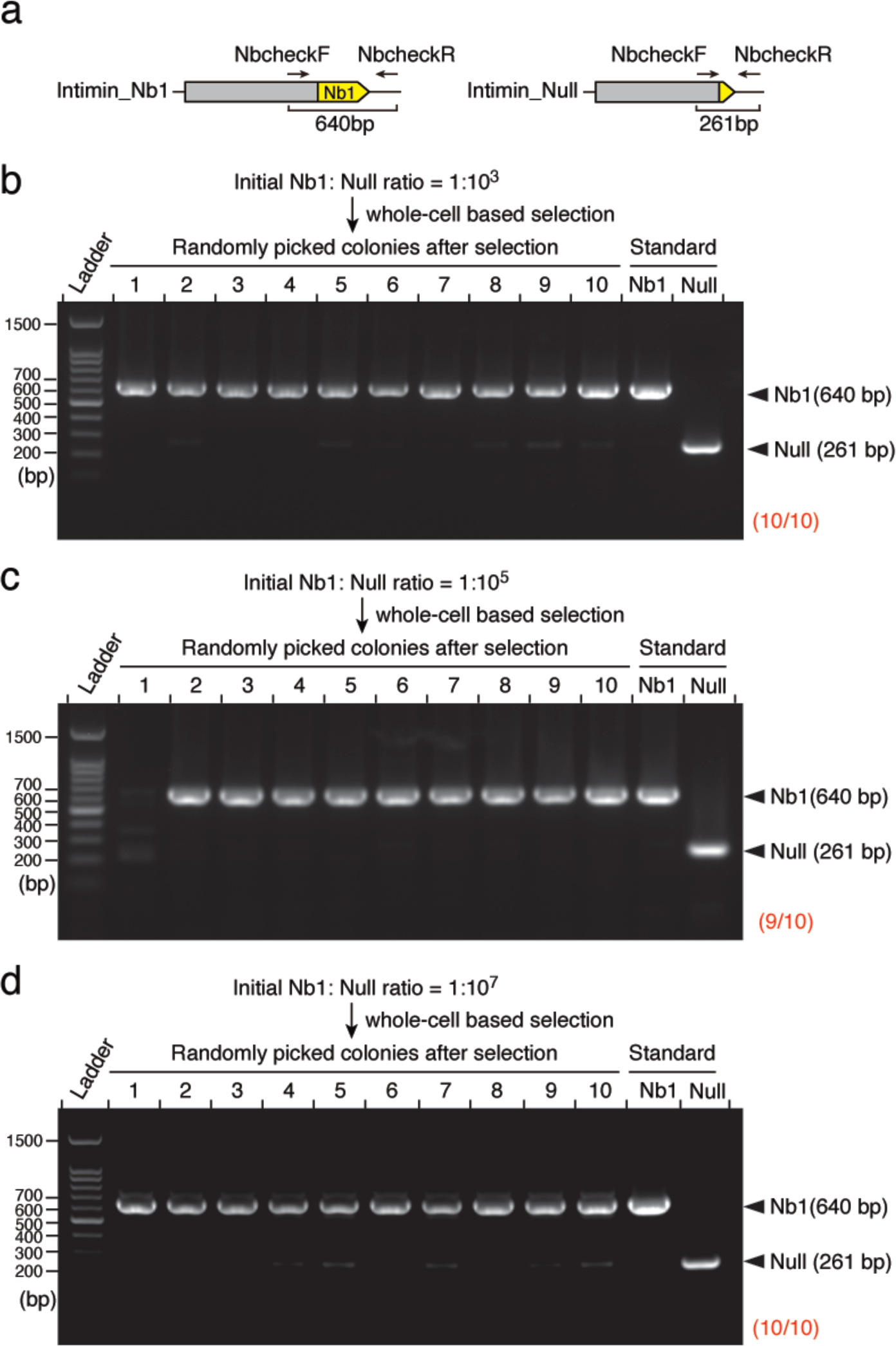
Diagnostic PCR verification of the efficacy of the selection strategy in enriching for low-abundance nanobodies. (**a**) The display system employed in this study comprises the intimin N-terminus (grey) fused with the adhesin region (yellow) at the C-terminus. Arrows indicate the annealing primers utilized for diagnostic PCR. The expected length of the PCR products for the cells displaying Nb1 (640 bp) and null (261 bp) are underscored. (**b**) PCR analysis of the transconjugants after selection. Two recipient strains expressing Nb1 and non-matched null control, respectively, were mixed at an initial ratio of 1:10^3^. The primers illustrated in (**a**) were used for diagnostic PCR. The proportions of colonies expressing Nb1 are denoted at the bottom-right (red). (**c**) Randomly selected colonies after two rounds of bio-panning were subjected to diagnostic PCR. The recipient cells displaying Nb1 were diluted with cells expressing a non-matched null control at an initial ratio of 1:10^5^. (**d**) The recipient population, initially at a ratio of Nb1:Null = 1:10^7^, after three rounds of selection was verified through diagnostic PCR.

**Fig. S3.**
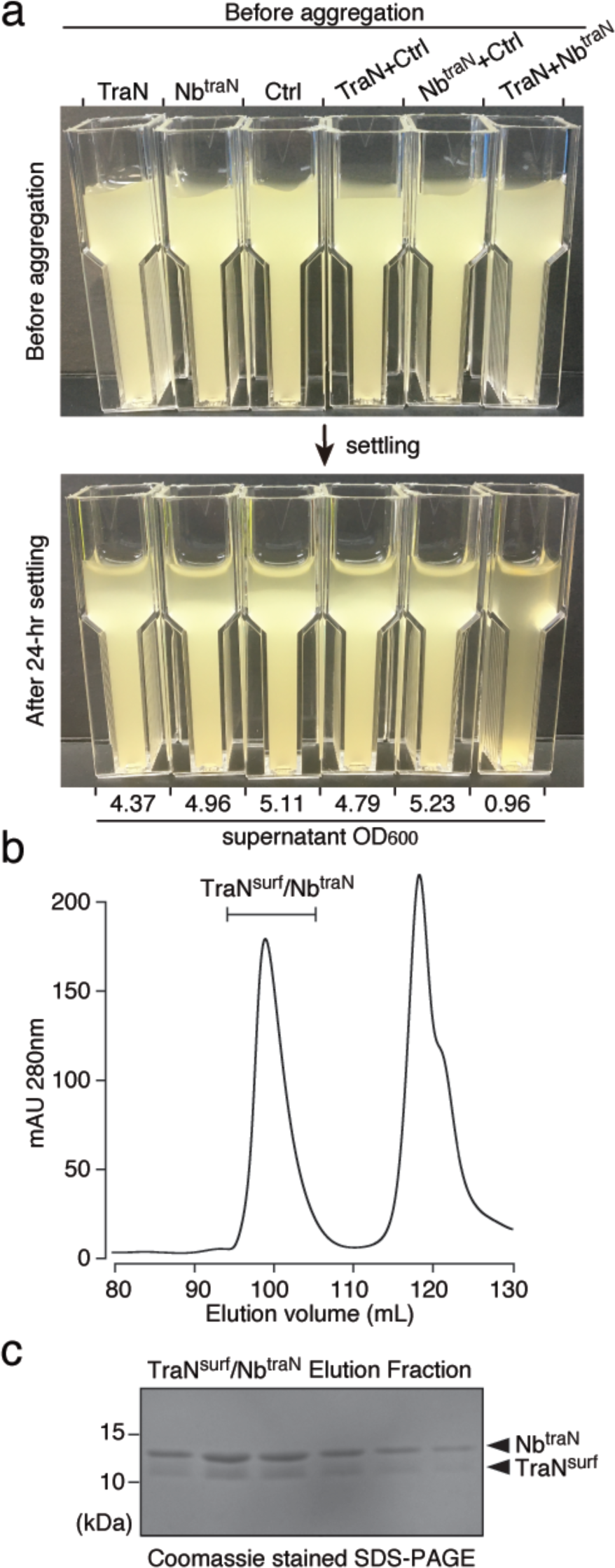
Nb^traN^ binds to TraN. (**a**) The macroscopic aggregation analysis to assess cell-cell adhesion for the indicated cultures of *E. coli* strains either individually or in combination. *E. coli* displaying TraN, Nb^traN^, or the non-matched control (Ctrl) were tested. The OD_600_ of the culture supernatants was measured after the settling period. (**b**) Size-exclusion chromatography profile of the Nb^TraN^/TraN^surf^ complex using a HiLoad 16/600 Superdex 200pg column. (**c**) The coomassie-stained SDS-PAGE protein gel shows the elution fractions of Nb^traN^ in complex with TraN^surf^. Migrations of a size standard, in kDa, are indicted (left).

**Fig. S4.**
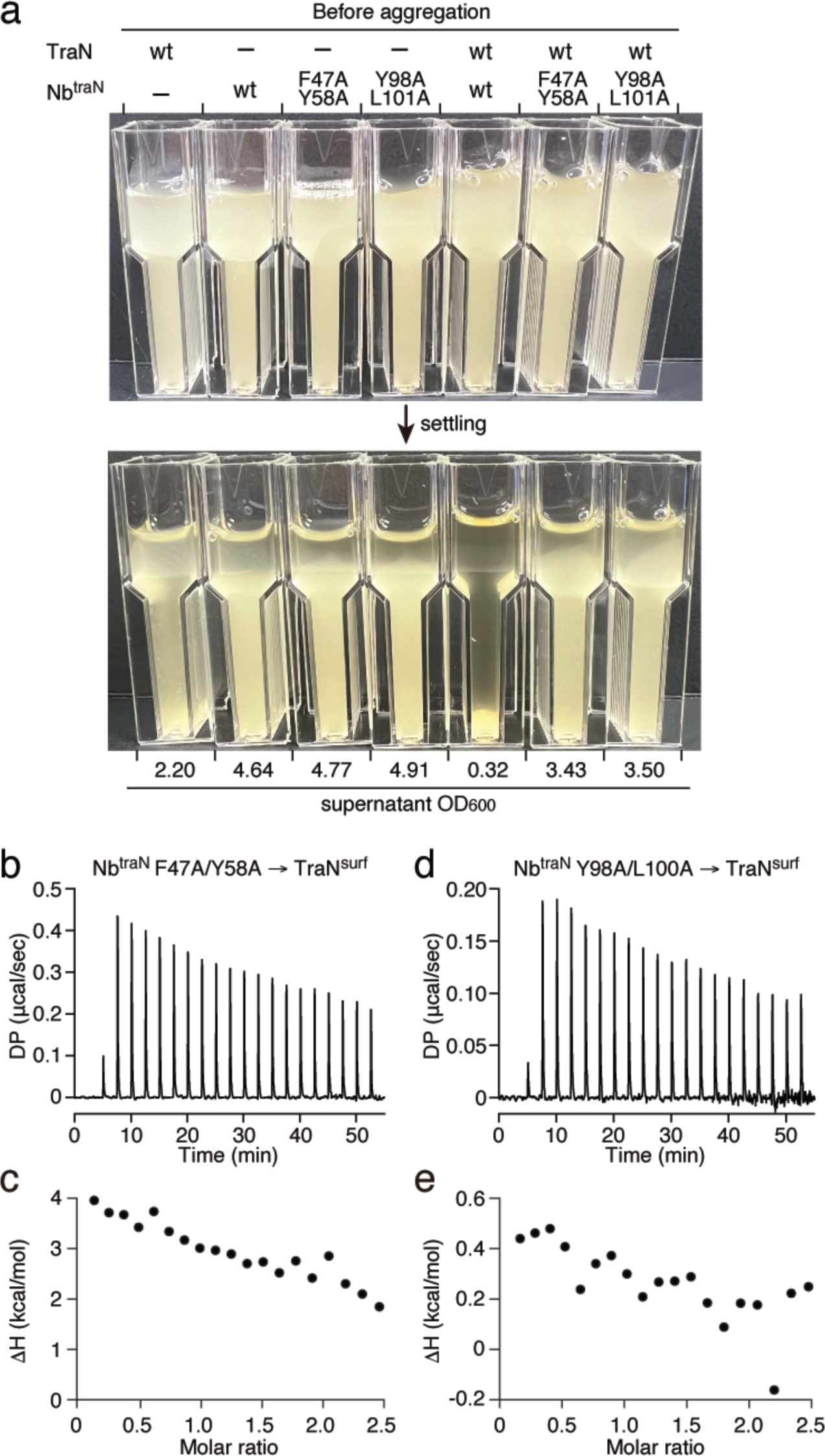
Mutations in the CDR2 or CDR3 of Nb^traN^ affect its interaction with TraN. (**a**) Comparison of the adhesive capability of the wild-type (wt) Nb^traN^ and its variants (CDR2: F47A/Y58A; CDR3: Y98A/L101A). For the macroscopic aggregation analysis, single cultures and mixtures were used. After settling, the OD_600_ of the culture supernatants was measured and values are shown below the image. (**b, c**) The ITC binding profiles of Nb^traN^ F47A/Y58A with TraN^surf^. (**d, e**) The ITC binding profiles of Nb^traN^ Y98A/L101A with TraN^surf^.

**Fig. S5.**
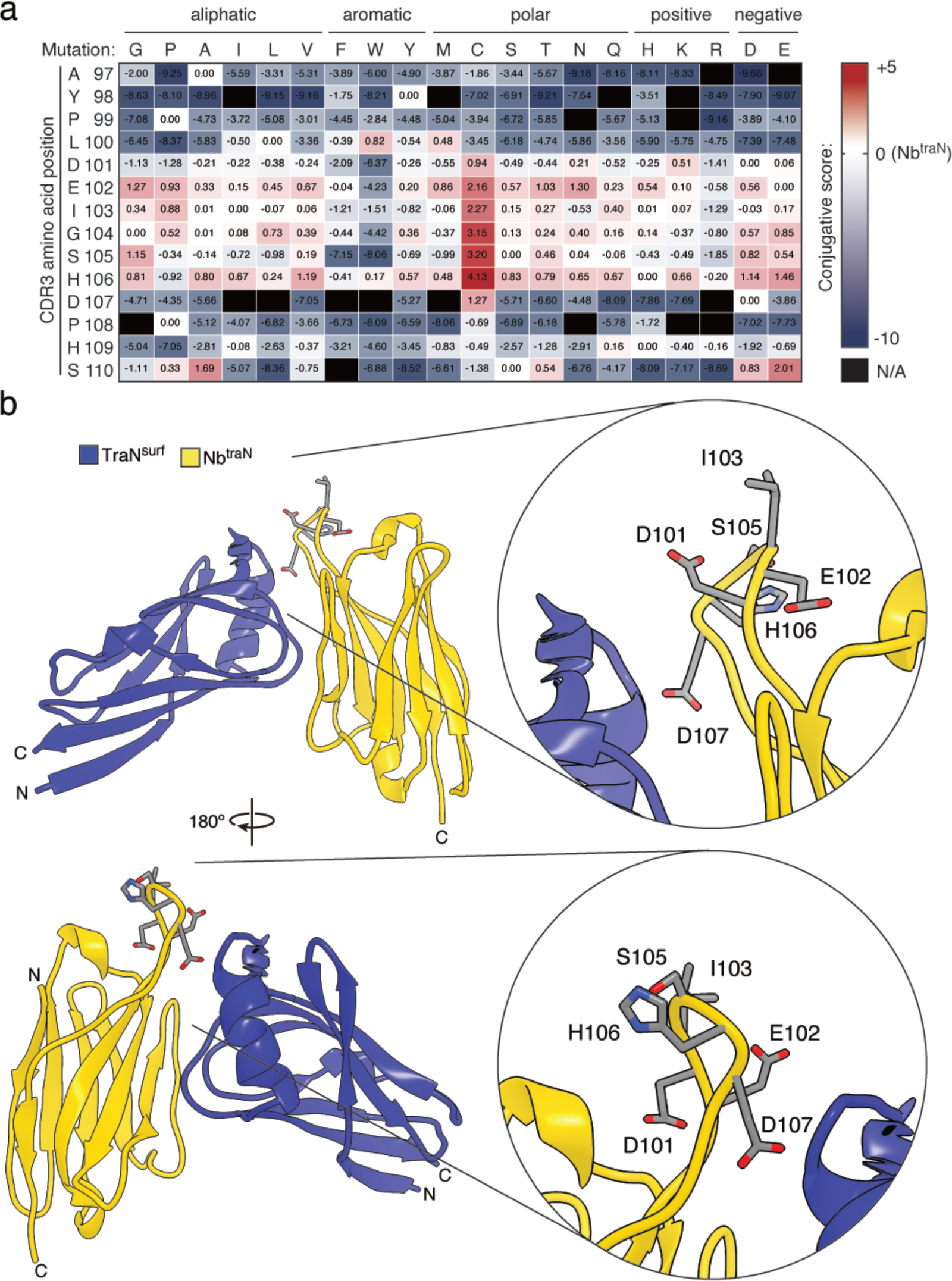
Deep mutational engineering of CDR3 in Nb^traN^. (**a**) The values of the conjugative scores of single amino acid substitution within CDR3 of Nb^traN^. Related to **Fig. 4**. (**b**) Crystal Structure of Nb^traN^ (yellow) bound to the surface-exposed domain of TraN (TraN^surf^, blue) (PDB: 8X7N). A second view (rotated 180° around the vertical axis) is shown below. The close-up views on the right side indicate the locations of the cysteine variants (residues 101 to 107) on Nb^traN^.

**Fig. S6.**
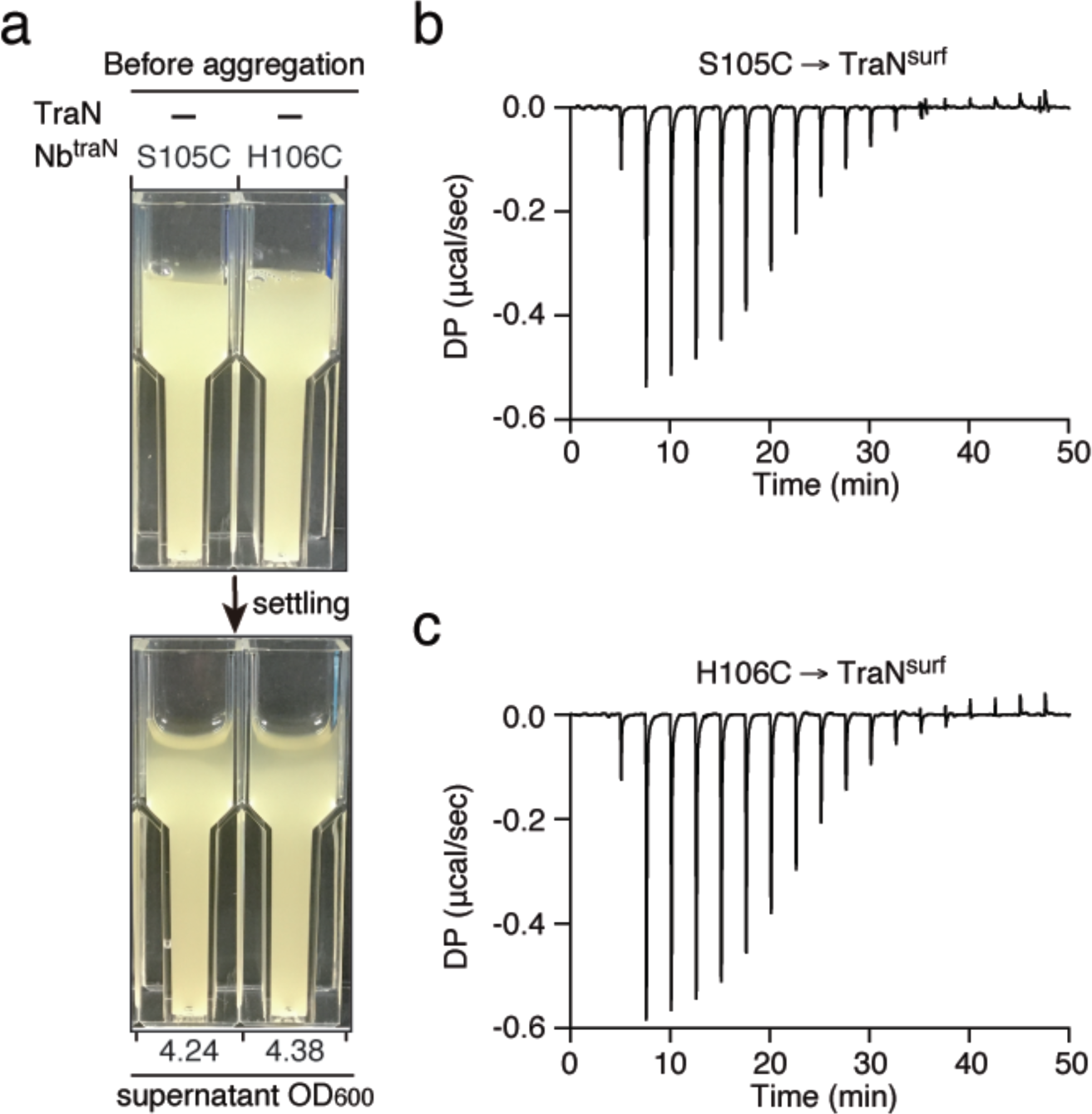
Residue substitutions of CDR3 in Nb^traN^ improve its binding to TraN. (**a**) Control cultures of *E. coli* strains expressing the Nb^traN^ variants that were used in the macroscopic aggregation assay in **Fig. 4**. (**b, c**) The ITC binding profiles of Nb^traN^ S105C (**b**) and H106C (**c**) with TraN^surf^. Related to **Fig. 4**.

**Fig. S7.**
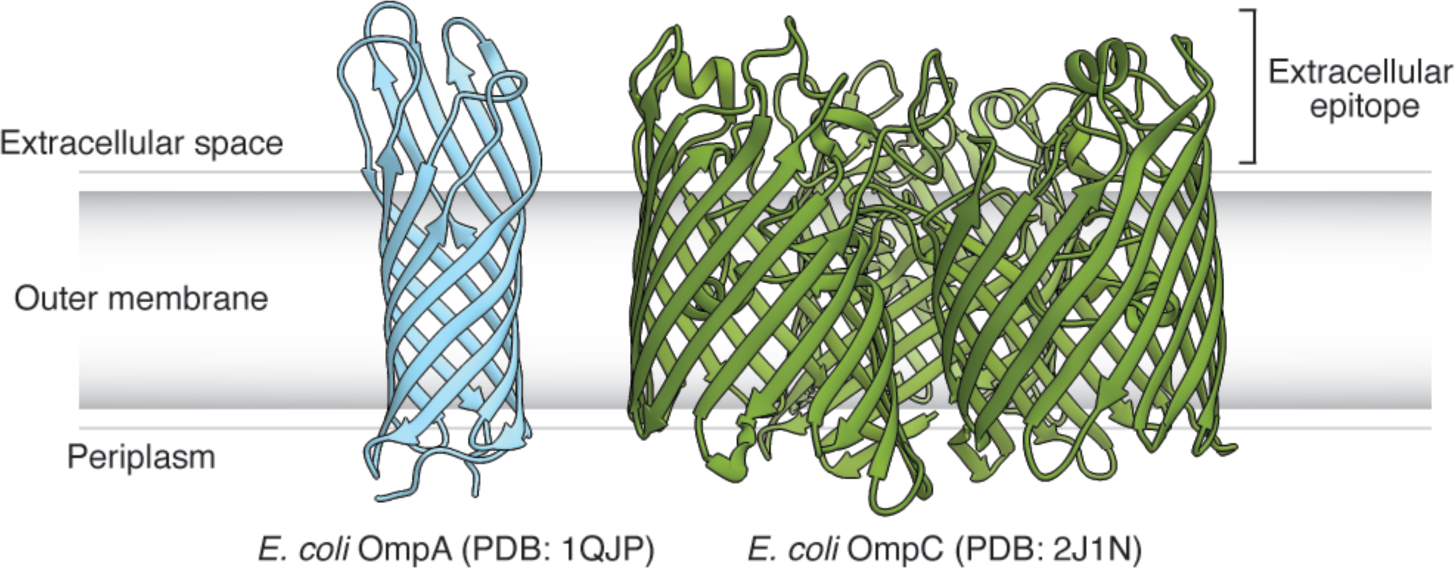
Structures of *E. coli* OmpA and OmpC. Crystal structures of the OmpA membrane domain (PBD: 1QJP)^1^ and OmpC (PDB: 2J1N)^2^ from *E. coli*. OmpA (blue) is presented as a monomeric structure, featuring an eight-stranded β-barrel spanning the outer membrane, coupled with flexible loops at its external end. Homotrimeric OmpC (green) is composed of three beta-barrels. The monomeric form of OmpC exhibits a 16-stranded β-barrel porin fold, with irregular extracellular loops.

**Fig. S8.**
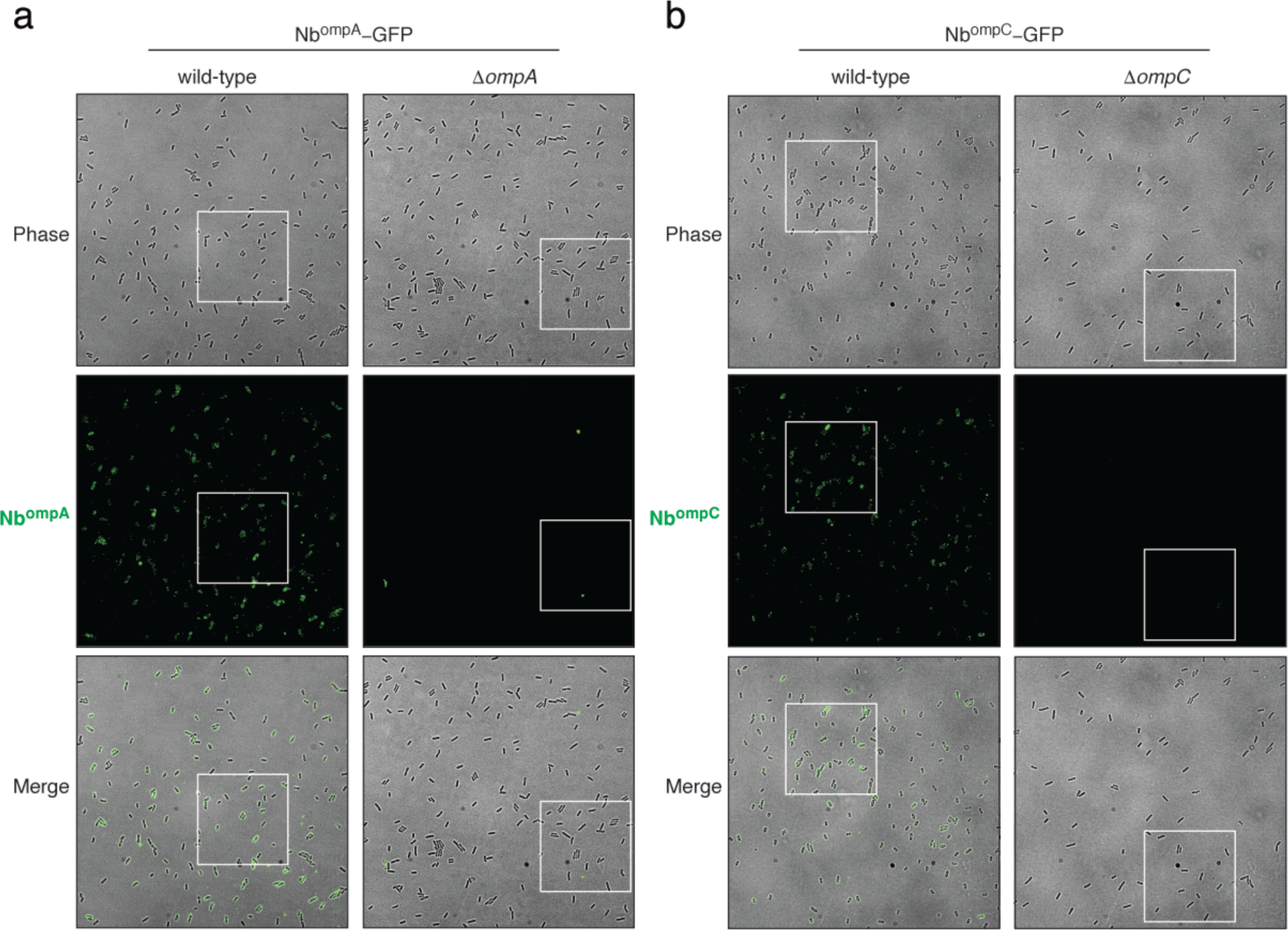
Nb^ompA^-GFP and Nb^ompC^-GFP bind to their target antigens. (**a**) The phase-contrast (top), green fluorescence (middle), and merged (bottom) images of *E. coli* (wild-type and *ΔompA* strain) following 1-hour incubation with Nb^ompA^-GFP. The white borders demarcate the cropped images displayed in **Fig. 5e**. (**b**) Fluorescence micrographs of *E. coli* (wild-type and *ΔompC* strain) cells treated with Nb^ompC^-GFP for 1 hour. Phase-contrast (top), green fluorescence (middle), and merged (bottom) images are presented. Zoomed-in images of the areas within white borders are featured in **Fig. 5f**.

**Fig. S9.**
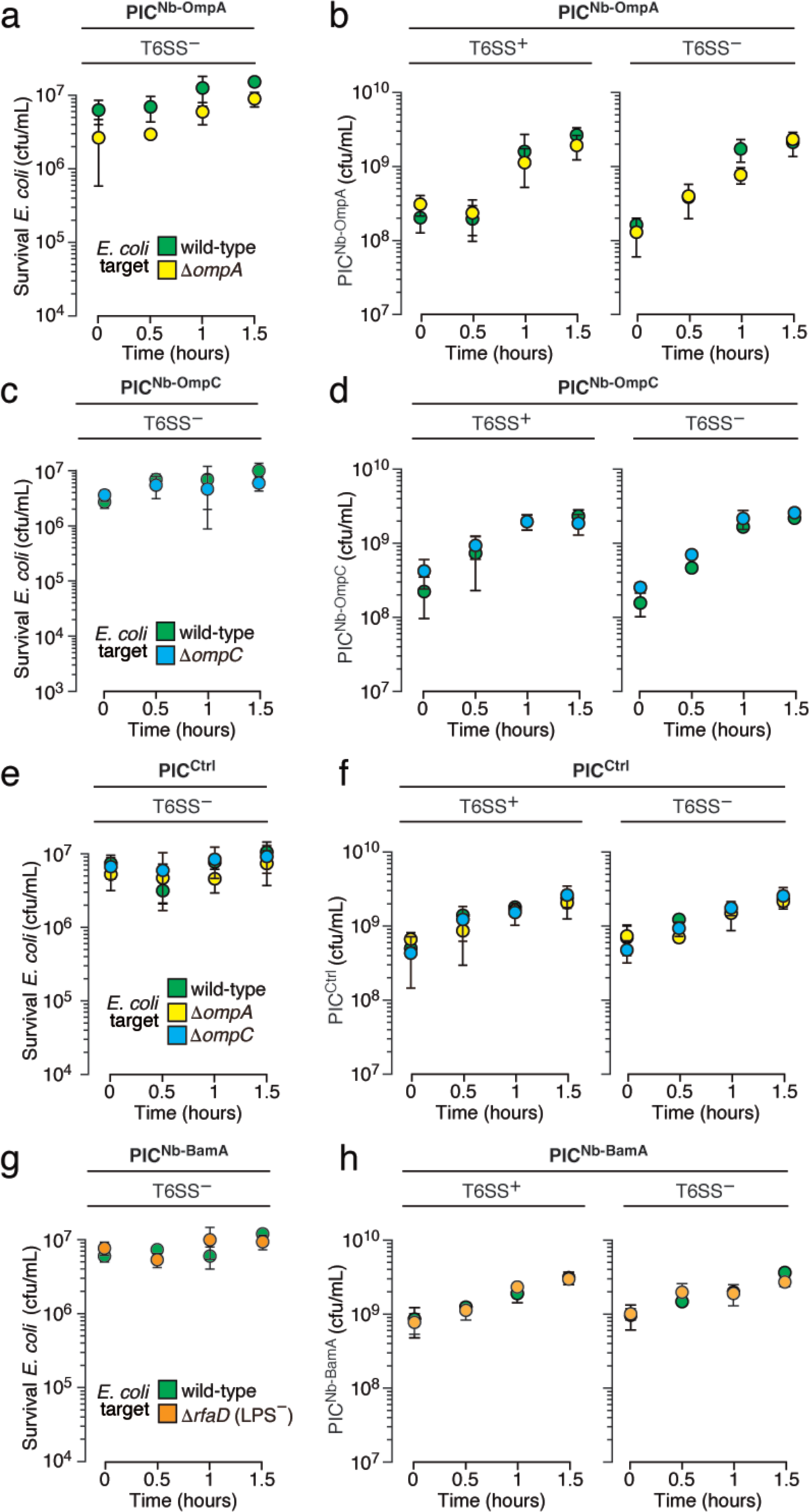
Target depletion mediated by PICs requires a functional T6SS, and survival of PICs upon incubation with target *E. coli*. (**a, c, e, g**) Survival (CFUs/mL) of *E. coli* MG1655 wild-type, Δ*ompA*, Δ*ompC*, and Δ*rfaD* strains (colored green, yellow, blue, and orange, respectively) upon liquid co-culture with T6SS-deficient PICs expressing Nb^ompA^ (**a**), Nb^ompC^ (**c**), non-matched control (**e**), or Nb^BamA^ (**g**). (**b, d, f, h**) The survival CFU of PICs expressing Nb^ompA^ (**b**), Nb^ompC^ (**d**), non-matching control (**f**), or Nb^BamA^ (**h**) upon liquid co-culture with target cells *E. coli* MG1655 wild-type (green), Δ*ompA* (yellow), Δ*ompC* (blue), and Δ*rfaD* (orange) cells. PICs with a functional (T6SS+) or dysfunctional T6SS (T6SS−, Δ*icmF*) were utilized. Data are presented as means ± SD. n = 3 technical replicates, representative of two biological replicates. *p < 0.05, t-test.

**Fig. S10.**
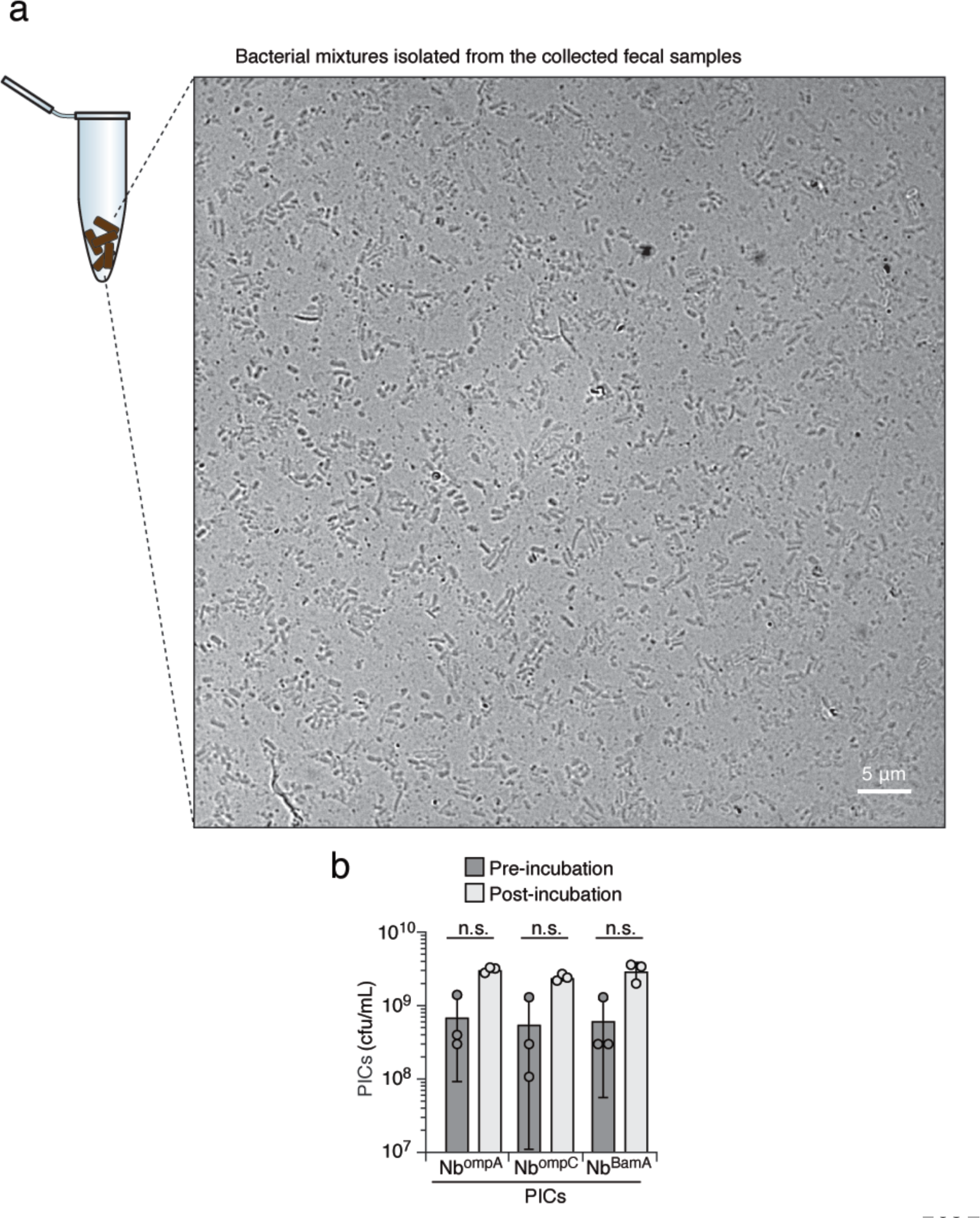
Survival of PICs remains unaffected before and after incubation in a bacterial mixture derived from mice fecal samples. **(a)** Phase-contrast micrograph of a bacterial population isolated from mouse fecal samples. Scale bar = 5 μm. (**b**) Survival (CFUs/mL) of PICs before and after incubation with mouse fecal microbiota. Related to **Fig. 6f**.

**Supplementary Table 1.** X-ray data collection and refinement statistics. Relevant to Method and Fig. 3.

| Crystals | TraN<sup>surf</sup>/Nb<sup>traN</sup> complex |
| --- | --- |
| Space group | 2<sub>1</sub> 2<sub>1</sub> 2<sub>1</sub> |
| Cell dimensions |  |
| a, b, c (Å) | a= 96.864<br>b=153.263<br>c=163.03 |
| α, β, γ (o) | α=β=γ=90 |
| Data collection |  |
| Wavelength (Å) | 0.97625 |
| Resolution | 19.85 -3.67 |
| Total reflections | 230515 |
| Unique reflections | 26580 |
| R<sub>merge</sub> (%)<sup>a</sup> | 20.5 (63.9) |
| CC1/2 (%)<sup>a</sup> | 98.8 (95.5) |
| I / σI<sup>a</sup> | 7.3 (2.6) |
| Completeness (%) | 99.8 (98.5) |
| Redundancy | 8.6 |
| Refinement |  |
| R<sub>work</sub> / R<sub>free</sub> (%) | 25.96/28.54 |
| Average B-factor | 84.2 |
| Bond length rmsd (Å) | 0.002 |
| Bond angle rmsd (°) | 0.478 |
| Ramachandran plot (%) |  |
| Favored region | 94.10 |
| Outlier region | 0.0 |
| PDB code | 8X7N |

<sup>a</sup> Statistics for the highest-resolution shell are shown in parentheses.

**Supplementary Table 2.**
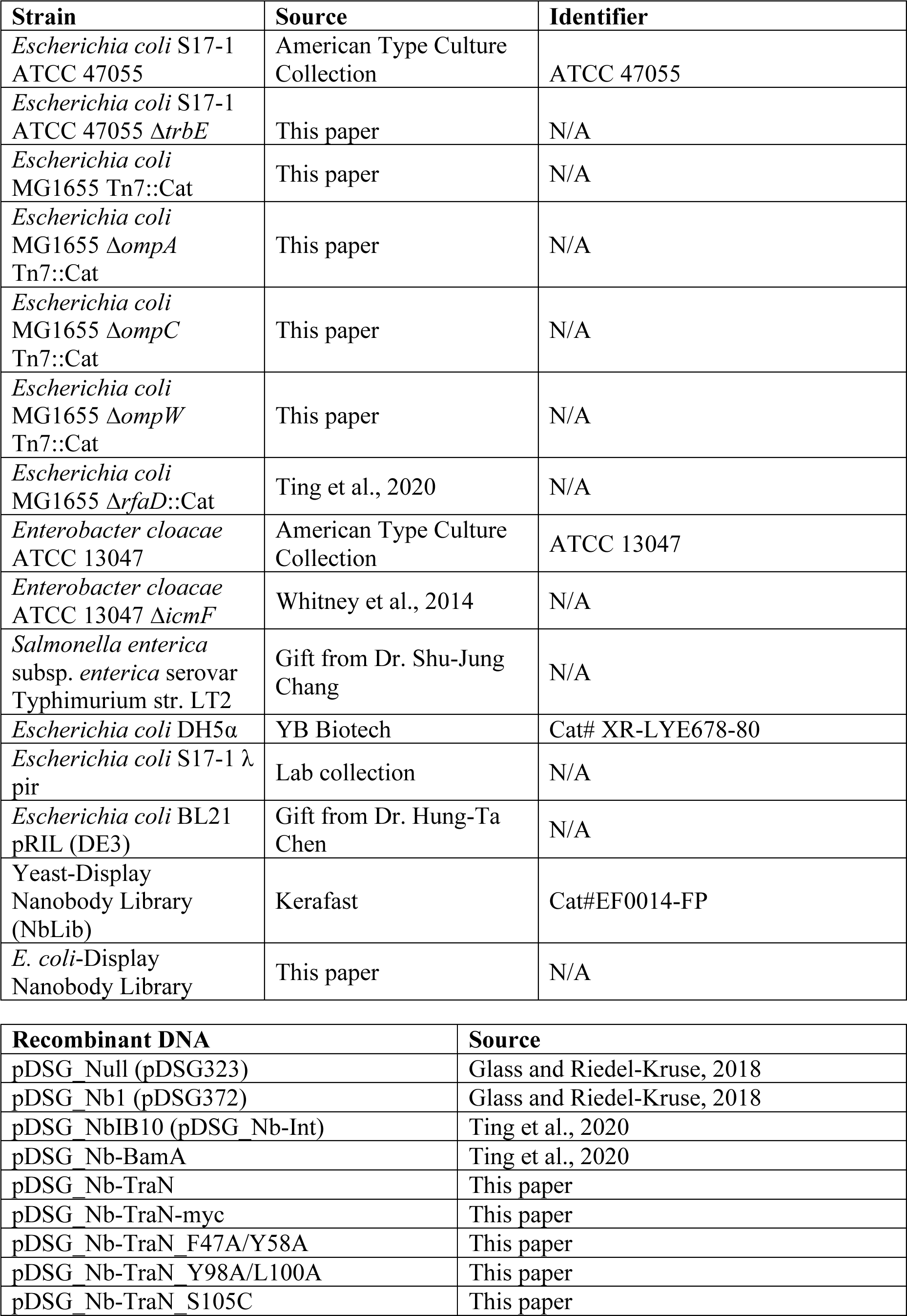
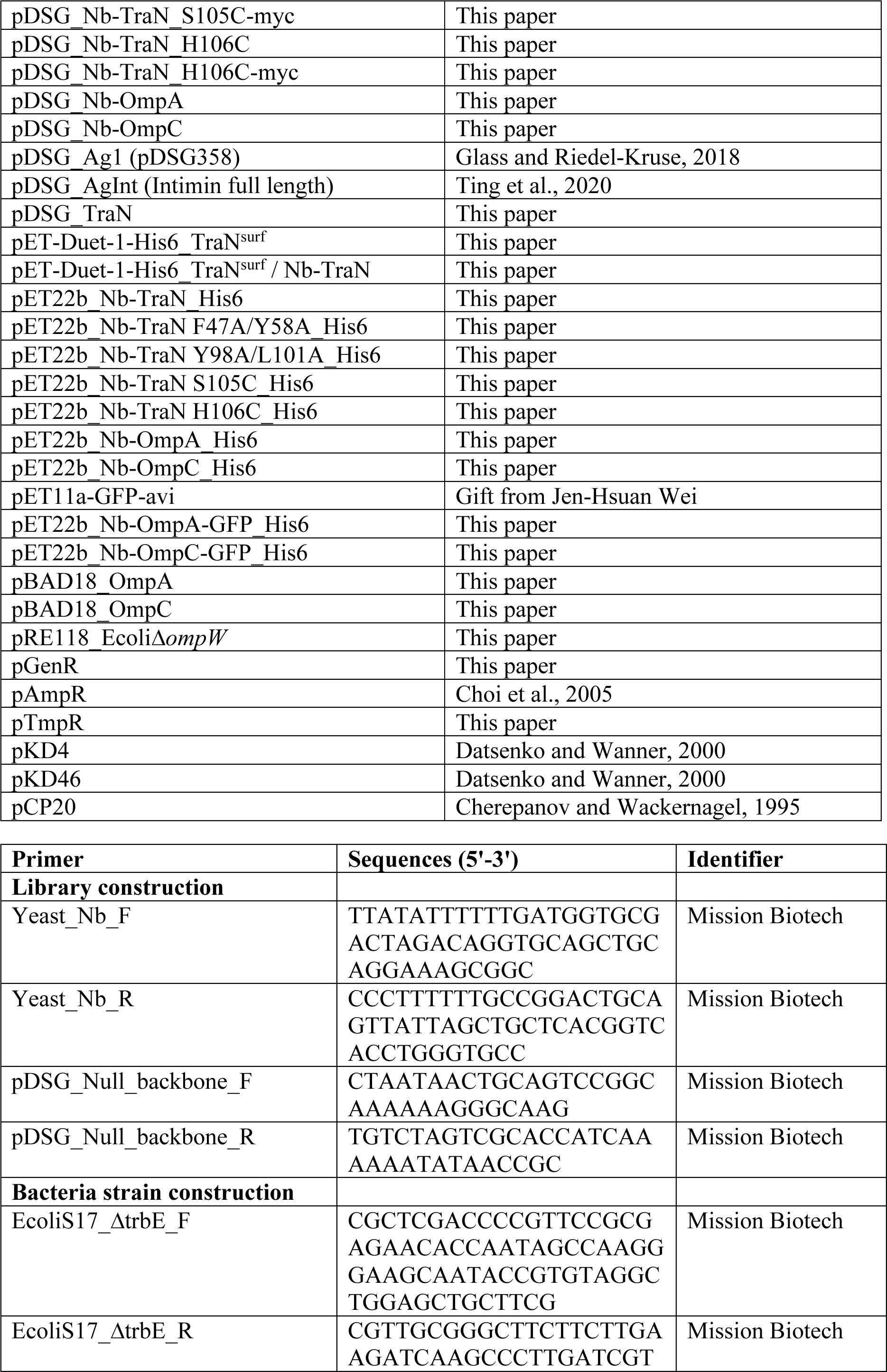
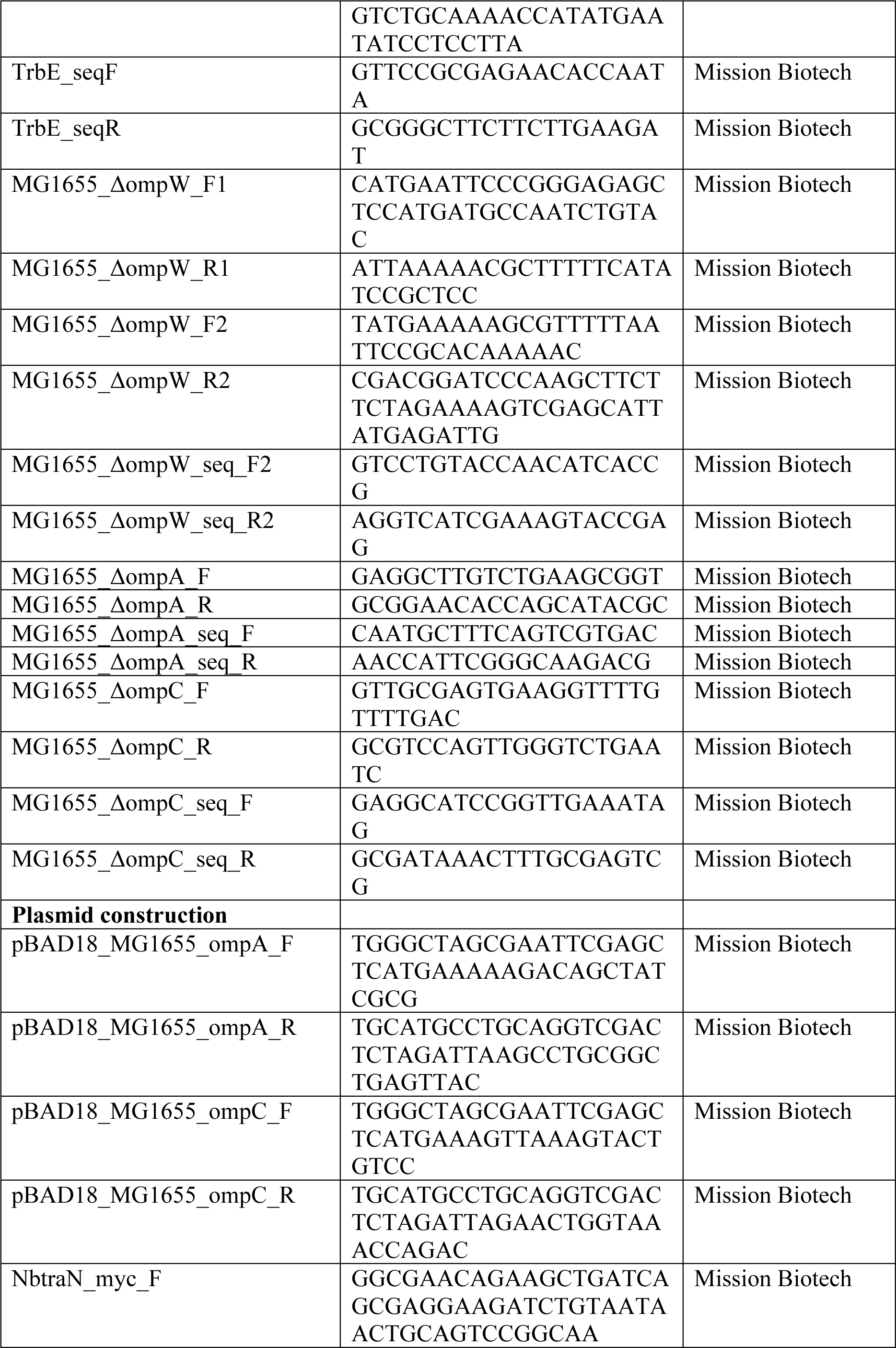
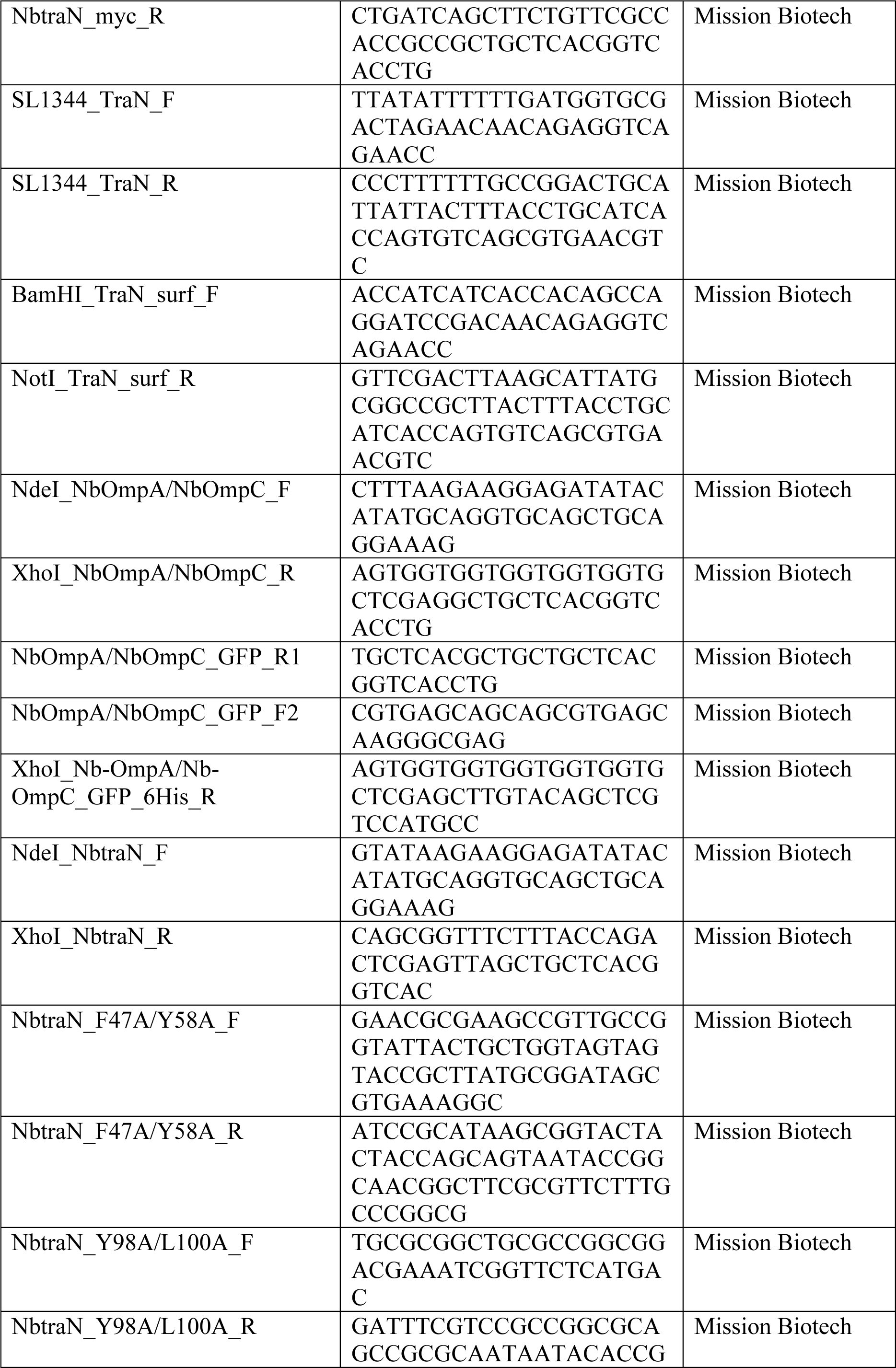
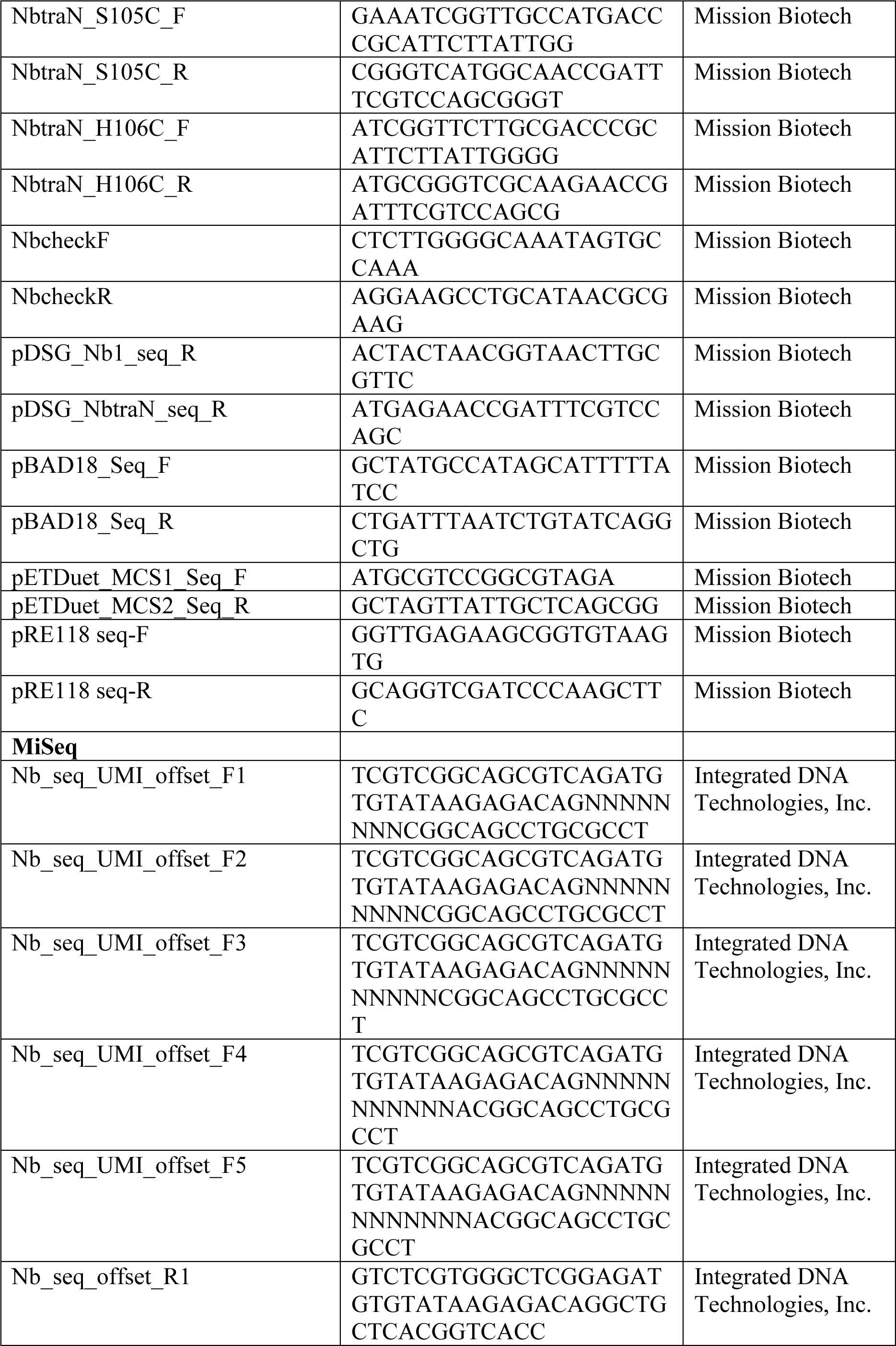
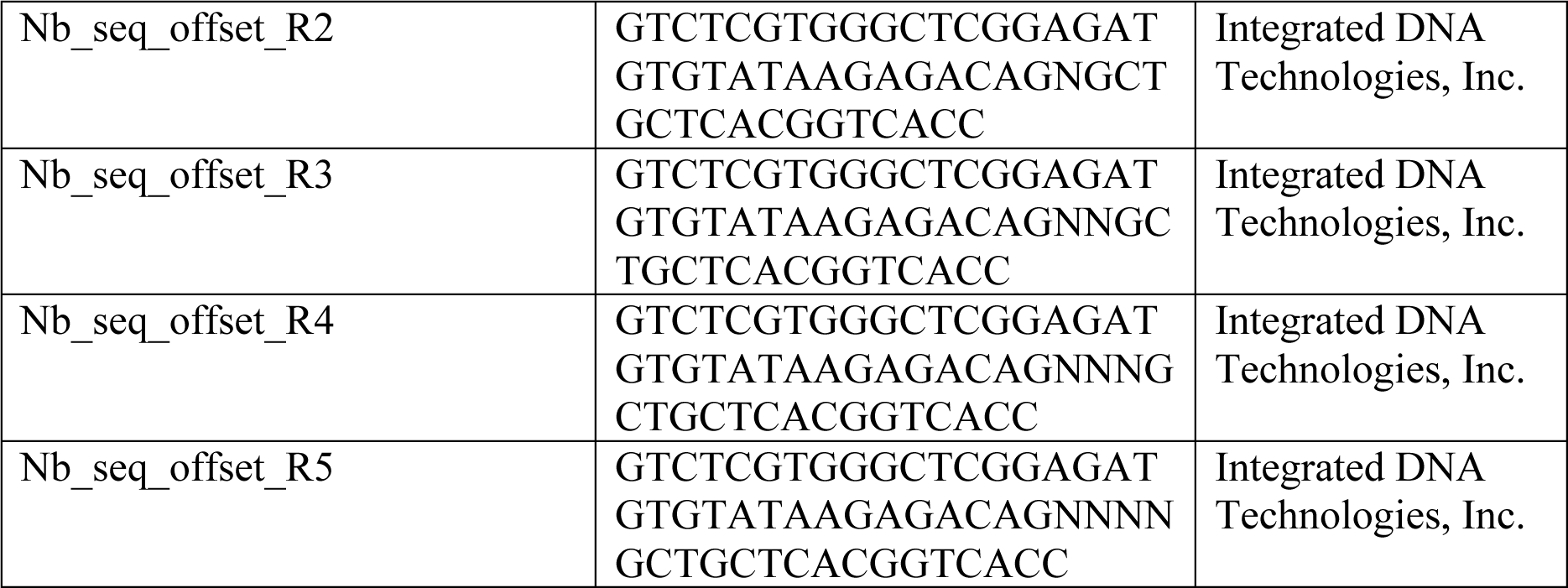
Strains, Recombinant DNA, and Oligonucleotides used in this study.

**Supplementary Table 3.** Illumina MiSeq deep-sequencing analysis of the nanobodies against TraN antigen. Related to Fig. 3 and 4.

## DATA AVAILABILITY

The coordinates and structure factors for the Nb^traN^/TraN^surf^ complex have been deposited in the Protein Data Bank, with accession code 8X7N. Sequence data associated with this study is available from the Sequence Read Archive at GSE249103.

## AUTHOR CONTRIBUTIONS

P.-Y.C., W.-L.W., K.-C.H., and S.-Y.T. designed the study. P.-Y.C., Y.-C.C., P.-P.C., K.-T.L., and K.-C.H. performed experiments. P.-Y.C., Y.-C.C., P.-P.C., K.-C.H., and S.-Y.T. analyzed data. P.-Y.C., Y.-C.C., P.-P.C., K.-C.H., and S.-Y.T. wrote the manuscript.

